# Predicting higher-order mutational effects in an RNA enzyme by machine learning of high-throughput experimental data

**DOI:** 10.1101/2022.05.31.494017

**Authors:** James D. Beck, Jessica M. Roberts, Joey Kitzhaber, Ashlyn Trapp, Edoardo Serra, Francesca Spezzano, Eric J. Hayden

## Abstract

Ribozymes are RNA molecules that catalyze biochemical reactions. Self-cleaving ribozymes are a common naturally occurring class of ribozymes that catalyze site-specific cleavage of their own phosphodiester backbone. In addition to their natural functions, self-cleaving ribozymes have been used to engineer control of gene expression because they can be designed to alter RNA processing and stability. However, the rational design of ribozyme activity remains challenging, and many ribozyme-based systems are engineered or improved by random mutagenesis and selection (in vitro evolution). Improving a ribozyme-based system often requires several mutations to achieve the desired function, but extensive pairwise and higher-order epistasis prevent a simple prediction of the effect of multiple mutations that is needed for rational design. Recently, high-throughput sequencing-based approaches have produced data sets on the effects of numerous mutations in different ribozymes (RNA fitness landscapes). Here we used such high-throughput experimental data from variants of the CPEB3 self-cleaving ribozyme to train a predictive model through machine learning approaches. We trained models using either a random forest or long short-term memory (LSTM) recurrent neural network approach. We found that models trained on a comprehensive set of pairwise mutant data could predict active sequences at higher mutational distances, but the correlation between predicted and experimentally observed self-cleavage activity decreased with increasing mutational distance. Adding sequences with increasingly higher numbers of mutations to the training data improved the correlation at increasing mutational distances. Systematically reducing the size of the training data set suggests that a wide distribution of ribozyme activity may be the key to accurate predictions. Because the model predictions are based only on sequence and activity data, the results demonstrate that this machine learning approach allows readily obtainable experimental data to be used for RNA design efforts even for RNA molecules with unknown structures. The accurate prediction of RNA functions will enable a more comprehensive understanding of RNA fitness landscapes for studying evolution and for guiding RNA-based engineering efforts.

## Introduction

RNA enzymes, or ribozymes, are structured RNA molecules that catalyze biochemical reactions. One well-studied class of ribozymes are the small self-cleaving ribozymes that catalyze site specific cleavage of phosphate bonds in their own RNA backbone (Ferré-D’Amaré and Scott, 2010). These self-cleaving ribozymes are found in all domains of life, and their biological roles are still being investigated (Jimenez et al., 2015). In addition to their natural functions, these ribozymes have been used as the basis for engineering biological systems. For example, several small ribozymes (hammerhead, twister, pistol and HDV) have been used as genetically encoded gene regulatory elements by combining them with RNA aptamer and embedding them into untranslated regions of genes (Groher and Suess, 2014; Dykstra et al., 2022). This approach continues to gain attention because of the central importance of controlling gene expression and the simple design and build cycles of these small RNA elements. Nevertheless, ribozymes often need optimization for sequence dependent and cell specific effects. This can be achieved by modifying the sequence of the ribozymes, but this often requires multiple mutational changes and the vast sequence space requires extensive trial and error. Given this large sequence space, even the most high-throughput approaches can only find the optimal solutions present in the sequences that can be explored experimentally, which is a fraction of the total possible sequences. The engineering of ribozyme-based systems could benefit from accurate prediction of the effects of multiple mutations in order to narrow the search space towards optimal collections of sequences.

One way to think of the ribozyme optimization problem is in terms of fitness landscapes. Molecular fitness landscapes of protein and RNA molecules are studied by measuring the effects of numerous mutations on the function of a given reference molecule (Athavale et al., 2014; Blanco et al., 2019). Recently, the fitness landscapes of RNA molecules have been studied experimentally by synthesizing large numbers of sequences and using high-throughput sequencing to evaluate the relative activity of the RNA in vitro, or the growth effect of the RNA in a cellular system, both of which are termed “RNA fitness” (Kobori and Yokobayashi, 2016; Li et al., 2016; Pressman et al., 2019). The goal of in vitro evolution is often to find the highest peak in the landscape, or one of many high peaks, by introducing random mutations and selecting for improved activity. However, the RNA fitness landscapes that have been experimentally studied so far have revealed rugged topographies with peaks of high relative activity and adjacent valleys of low activity. Landscape ruggedness is an impediment to finding desired sequences through in vitro evolution approaches (Ferretti et al., 2018). Epistasis, defined as the non-additive effects of mutations, is the cause of ruggedness in fitness landscapes, and epistasis has been used to quantify the ruggedness of fitness landscapes (Szendro et al., 2013). More frequent and more extreme epistasis indicates that a landscape is more rugged. Importantly, more epistasis also means that the effect of combining multiple mutations is challenging to predict even if the effects of each individual mutation are known. In addition, experimental fitness landscapes can only study a limited number of sequences, except for very small RNA molecules (Pressman et al., 2019). It is often not possible to know if the process of in vitro evolution discovered a sequence that is globally optimal, or just a local optimum. For these reasons, it has become a goal to accurately predict the activity of sequences in order to streamline RNA evolution experiments and to study fitness landscapes in a more comprehensive manner (Groher et al., 2019; Schmidt and Smolke, 2021).

Here, we use high-throughput experimental data of mutational variants of a self-cleaving ribozyme to train a model for predicting the effect of higher-order combinations of three or more mutations. The ribozyme used in this study is the CPEB3 ribozyme (Figure 1A). This ribozyme is highly conserved in the genomes of mammals, where it is found in an intron of the CPEB3 gene (Salehi-Ashtiani et al., 2006). For training purposes, we generated a new data set that includes all possible individual and pairs of mutations to the reference CPEB3 ribozyme sequence (Figure 1B). These mutations were made by randomization of the CPEB3 ribozyme sequence with a 3% per nucleotide mutation rate during chemical synthesis of the DNA template. We reasoned that given the extensive amount of pairwise epistasis in RNA (Bendixsen et al., 2017), this data set might be sufficient for predicting higher-order mutants. In addition, we used a second, previously published data set that included 27,647 sequences comprised of random permutations of mutations found in mammals that include up to 13 mutational differences from the same reference ribozyme (Bendixsen et al., 2021). This second data set not only contains higher-order mutational combinations, but also a broad range of self-cleaving activity (Figure 1D). In both data sets, the relative activity of each sequence was determined by the deep sequencing of co-transcriptional self-cleavage data, as previously described. Briefly, the mutated DNA template was transcribed in vitro with T7 RNA polymerase. The transcripts were prepared for Illumina sequencing by reverse transcription and PCR. Relative activity was determined as the *fraction cleaved*, defined by the fraction of sequencing reads that mapped to a specific sequence variant in the shorter, cleaved form relative to the total number of reads for that sequence variant.

**Figure 1.**
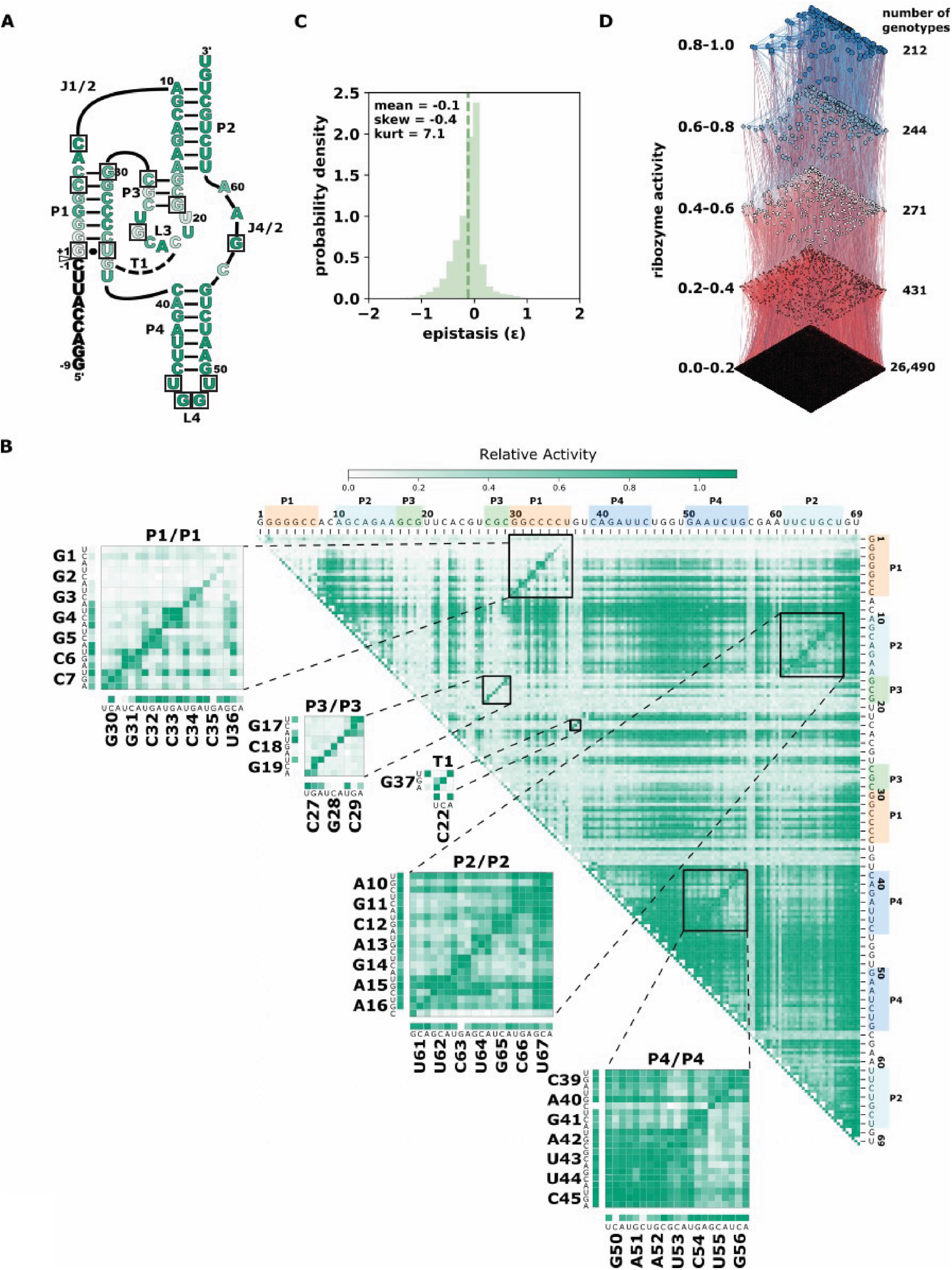
The CPEB3 ribozyme and data prediction challenge. (A) Secondary structure diagram of the CPEB3 ribozyme. The white arrow indicates the site of self-cleavage. Nucleotide color indicates the average relative activity of the three possible point mutations at each position. Boxes indicate nucleotide positions mutated in the phylogenetically derived higher-order mutants (B) Heatmap representation of comprehensive single and double-mutant data. Each pixel in the heatmap shows the ribozyme activity for a specific double mutant indicated by the nucleotide positions on the top and right of the heatmap. Insets show base paired regions and specific mutations. Ribozyme activity is determined as the fraction of total reads that map to each sequence that are in the cleaved form (fraction cleaved) relative to the wildtype fraction cleaved. (C) Distribution of pairwise epistasis from double mutant data. Epistasis was calculated as *ε =* log_10_ (*W*_AB_\**W*_wt_ / *W*_A_\**W*_B_), where *W*_wt_ is the fraction cleaved of the wild-type ribozyme, *W*_A_ and *W*_B_ are fraction cleaved of sequences with individual mutations and *W*_AB_ is the fraction cleaved of the sequence with both individual mutations. (D) Higher mutational distance variants of the CPEB3 ribozyme represented as a fitness landscape. *Ribozyme activity* (fraction cleaved) is shown for 27,647 sequence variants derived from permutations of naturally occurring mutations. Each node represents a different sequence and the size and color of the node is scaled to the ribozyme activity. Edges connect nodes that differ by a single mutation. Sequences are binned into quintiles of ribozyme activity and the *number of genotypes* reports the number of sequences in each quintile.

We set the goal of being able to predict the activity of the higher-order mutants in the phylogenetically derived fitness landscape (Figure 1D). In addition, we wanted to guide future experiments aimed at producing additional data for training models of ribozyme-based systems. The number of possible sequences increases exponentially with the number of variable nucleotide positions. In addition, the probability of finding active ribozymes at higher mutational distances becomes increasingly unlikely. Experiments aimed at training predictive models will need to choose realistic numbers of sequences that can have the highest impact on model performance. We therefore evaluated the effect of adding to the training data sequences with increasing mutational distances from the wild-type sequence as well as the effect of reducing the number of sequences in the training data. The results of these experiments were expected to be useful in guiding the choice of which sequence variants, and how many, to analyze experimentally in order to produce effective training data sets.

## Results

We first evaluated our new training data set that contained all single and double-mutants of the CPEB3 ribozyme. We found that the data did in fact contain full coverage of the possible 207 single mutants and the 21,114 double mutants. While the number of reads that mapped to each of these sequences varied, we found that, on average, 170 reads mapped to each double mutant, and ∼18,000 reads mapped to each single mutant (Supplemental Figure 1). This read depth was sufficient for the determination of the fraction cleaved for all single and double mutants (Figure 1B). Mapping the fraction cleaved to base paired structural elements showed expected patterns of activity caused by compensatory base pairs. Mutations that break a base pair typically showed low activity, but a second mutation that restored the base pair showed high activity. To further evaluate this data, we calculated the non-additive pairwise epistasis in this data set (Figure 1C). Together, this analysis indicated that this data set contained a wide range of ribozyme activity and the effects of all pairwise intramolecular epistatic interactions.

In order to determine the training potential of the comprehensive double-mutant data, we first trained models using only the fraction cleaved data for sequences with two or fewer mutations including the wild-type reference sequence. We then tested the models’ performance in predicting the fraction cleaved for sequences with increasing numbers of mutations. We trained two models with two approaches (see Materials and Methods). The first approach used a Random Forest regressor. In the second approach, we added a Long Short-Term Memory (LSTM) recurrent neural network to extract hidden features from the data. We then fed the hidden features with associated fraction cleaved to a Random Forest regressor. We will refer to this approach as “LSTM”. We found that models trained on 2 or fewer mutations with Random Forest outperformed LSTM at predicting the activity of sequences with five or fewer mutations (Figure 2 A-C), but LSTM performed better when predicting the activity of sequences with six or more mutations relative to the wild-type (Figure 2 D-I). However, both approaches showed a decrease in the correlation between predicted and observed when challenged to predict the activity of sequences with higher numbers of mutations, and both resulted in relatively low correlation (Pearson r < 0.7) for sequences with seven or more mutations when trained only on this double mutant data (Figure 2 and Supplementary Table 2). We concluded that models trained on simple random mutagenesis containing all double mutants can be useful for predicting lower mutational distances, but we anticipated that additional data might improve the ability to predict the effect of higher numbers of mutations.

**Figure 2.**
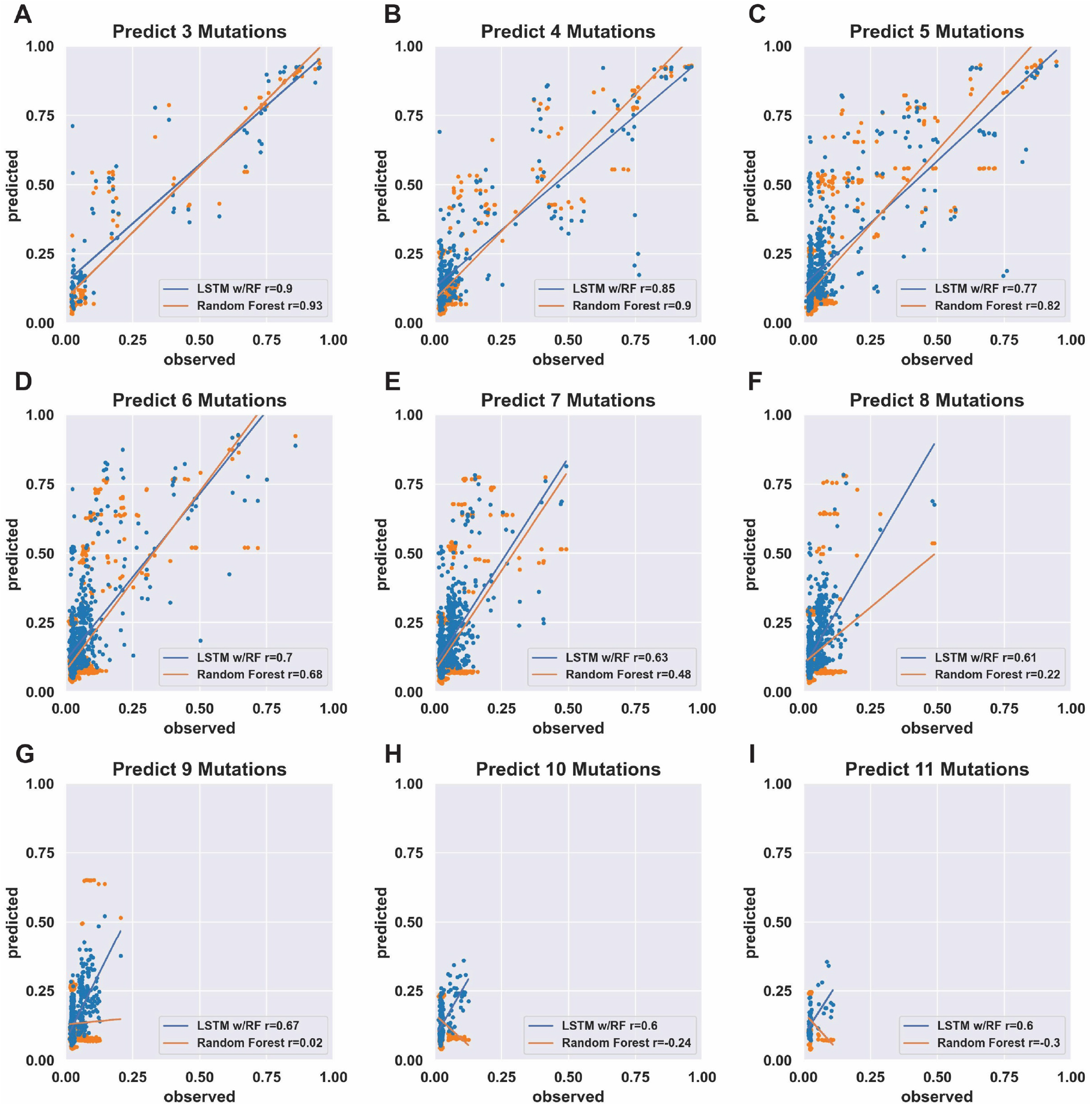
Prediction accuracy of models trained on comprehensive individual and pairs of mutations. (A-I) Scatter plots of *Predicted* (fraction cleaved from the models) and *Observed* (fraction cleaved from experiments). The models were trained on the experimentally determined fraction cleaved for the wild-type and all possible sequences with one mutation (207 sequences) or two mutations (21,114 sequences). Insets report Pearson correlation coefficients *r* for the model trained by the Random Forest approach (orange) and the LSTM-RF approach (blue). The sequences used to compare prediction vs. observed were separated by the number of mutations relative to the wild-type, as indicated by the title of each graph.

To determine the effect of adding higher-order mutants to the training data, we divided the phylogenetic derived sequence data by mutational distance and re-trained models with increasing orders of mutations in the training set. As expected, adding higher-order mutants improved the predicted to observed correlation at higher mutational distances (Figure 3 and Supplemental Figures 2-14). Interestingly, we found that the Random Forest approach outperformed the LSTM approach when sequences with more mutations were included in the training data. This is especially apparent for predicting the activity of sequences with 8-10 mutations. The Random Forrest approach resulted in models with high correlation between predicted and observed for all mutational distances when trained with data from sequences with four or more mutations (Figure 3 A-C). For both approaches, the largest improvements in the correlations occurred when sequences with 3 mutations (relative to wild-type) were added to the data. Subsequently appending additional sequences with greater numbers of mutations had diminishing improvements on the correlation. We note that all the testing data was set aside prior to training and identical testing data was used for all models. The results demonstrate that adding higher order mutants to the training data improves the Pearson correlation of sequences at higher distances in this data set. It is important to note that the phylogenetically derived data has different numbers of sequences for each class of mutations (Table 1), and sequences with higher numbers of mutations in our data show mostly low activity (Supplementary Figure 15). This helps interpret the effect of sequentially adding higher-order mutant sequences to the training data. It is also important to note that the phylogenetic derived sequences only contain mutations at thirteen different positions. The higher order sequences in this data are therefore combinations of the lower order sequences. For example, a sequence with six mutations can be constructed by combining two sequences with three mutations, both of which would be in the “3 mutations” training data. Our model is therefore predicting the effects of combining sets of mutations, and adding precise sets of lower order mutations that re-occur in higher order mutations clearly improves the correlations between prediction and experimental observation in our data.

**Figure 3.**
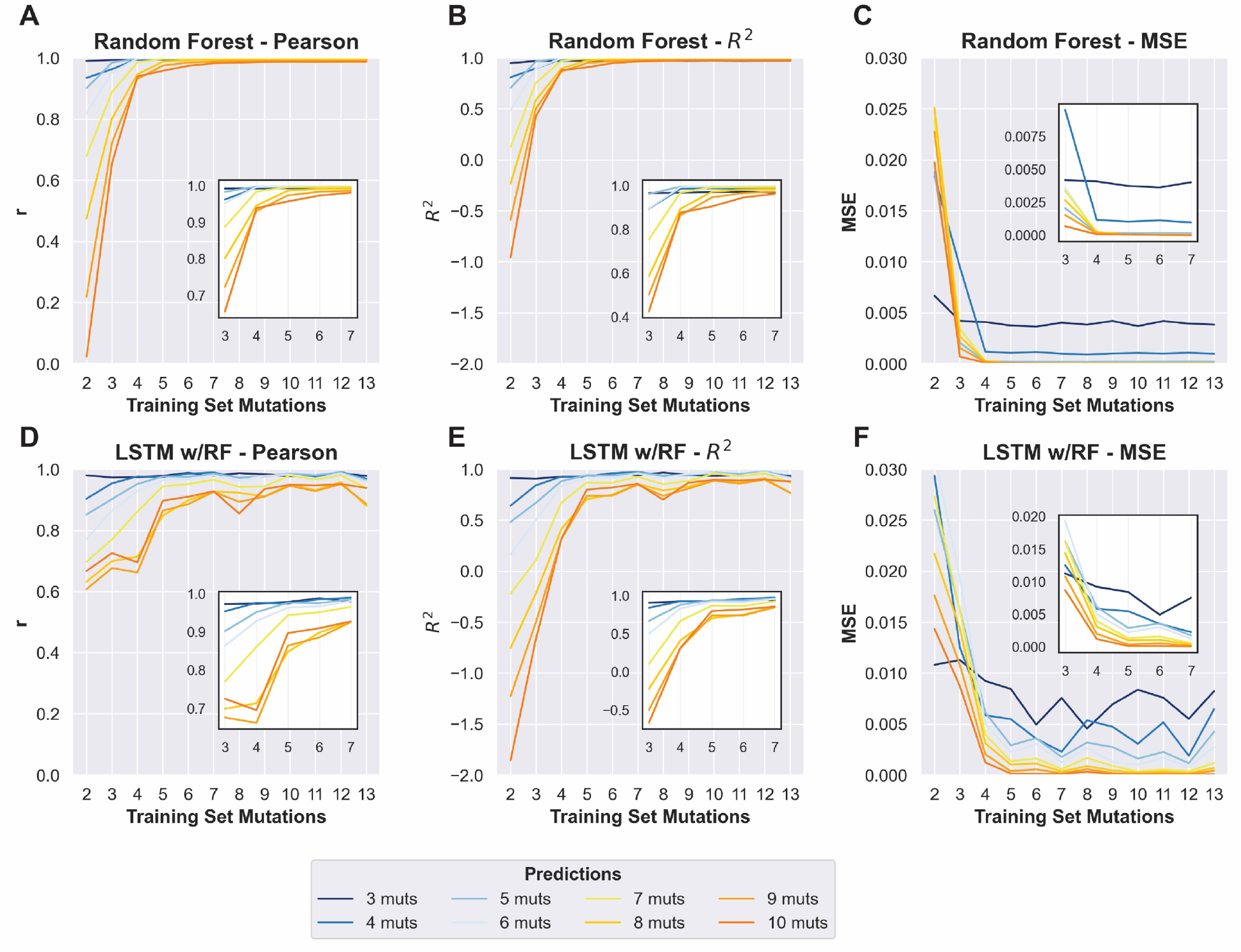
Improvement in prediction accuracy when including sequences with increased mutational distances in the training data. Changes in Pearson *r, R*^*2*^, and mean squared error (MSE) of prediction-observed correlation (y-axis) with increasing numbers of max mutations within the training data (x-axis). Training sets included all sequences up to and including the y-axis value. (A-C) Results obtained for the random forest model. (D-F) Results from the LSTM model. For each plot, colors indicate the numbers of mutations in sequences in the test data (see key). Insets show changes to the same prediction accuracy measurement with the 3-7 mutation training data, to allow more visual resolution.

**Table 1.** Counts of sequences in training and testing data sets.

| No. of mutations | Training | Testing |
| --- | --- | --- |
| 1 | 207 | --- |
| 2 | 21114 | --- |
| 3 | 414 | 104 |
| 4 | 1240 | 310 |
| 5 | 2650 | 662 |
| 6 | 4162 | 1040 |
| 7 | 4867 | 1217 |
| 8 | 4241 | 1060 |
| 9 | 2720 | 680 |
| 10 | 1249 | 312 |
| 11 | 389 | 97 |
| 12 | 74 | 18 |
| 13 | 6 | 2 |

In order to inform future experiments for collecting training data, we next set out to determine the effect of decreasing the amount of data in the training sets. Starting from the 80% of data used as prior training data, we randomly sampled sequences from this data to create new training data sets with 60%, 40%, 20%, 10% and 1% of the total data. These subsampled data sets were used to train models using the random forest regressor. The same testing data was set aside for all models and used to compare the Pearson correlation coefficient of each model trained with decreasing amounts of data. As an illustrative example, we focused on a model trained with sequences with 5 or fewer mutations relative to wild-type used to predict the activity of sequences with 7 mutations (Figure 4 and Supplementary Table 1). We chose this example because it achieved very high correlation (Pearson *r* = 0.99) when trained with 80% (25,733 unique sequences) of the data and therefore provided an opportunity to observe how rapidly the correlation decreased with less data. We found that the models trained on 5 or fewer mutations predicted with high correlation when as little as 40% (12,866) of the data was used for training (Pearson *r* = 0.97). With only 20% (6,433) and 10% (3,217) of the data, the model still showed good prediction accuracy with a Pearson correlation *r* ≅ 0.9. Surprisingly, we still observed reasonably high correlation when including only 1% (322) of the training data, and this was reproducible over five different models trained with different random samples of the data (Pearson *r* = 0.81, *stdev* = 0.046, *n = 5*). Similar results were observed with other training and testing scenarios. To illustrate general trends, we have plotted the Pearson correlation for the same model trained on 5 or fewer mutations when predicting the activity of sequences with 6, 7, 8 or 9 mutations, and for a model trained on 9 or fewer mutations used to predict sequences with 5, 6, 7, or 8 mutations (Figure 4). This analysis suggests that the total amount of training data is not critical for predicting the activity of sequences in our data set. When combined with the diminishing returns of adding more higher order mutations (Figure 3), this analysis emphasizes the importance of collecting appropriate experimental data sets for training that include ribozymes with more mutations that still maintain relatively high activity. However, given the low probability of finding higher-order sequences with higher activity, an iterative approach with several cycles of predicting and testing might be necessary to acquire such data.

**Figure 4.**
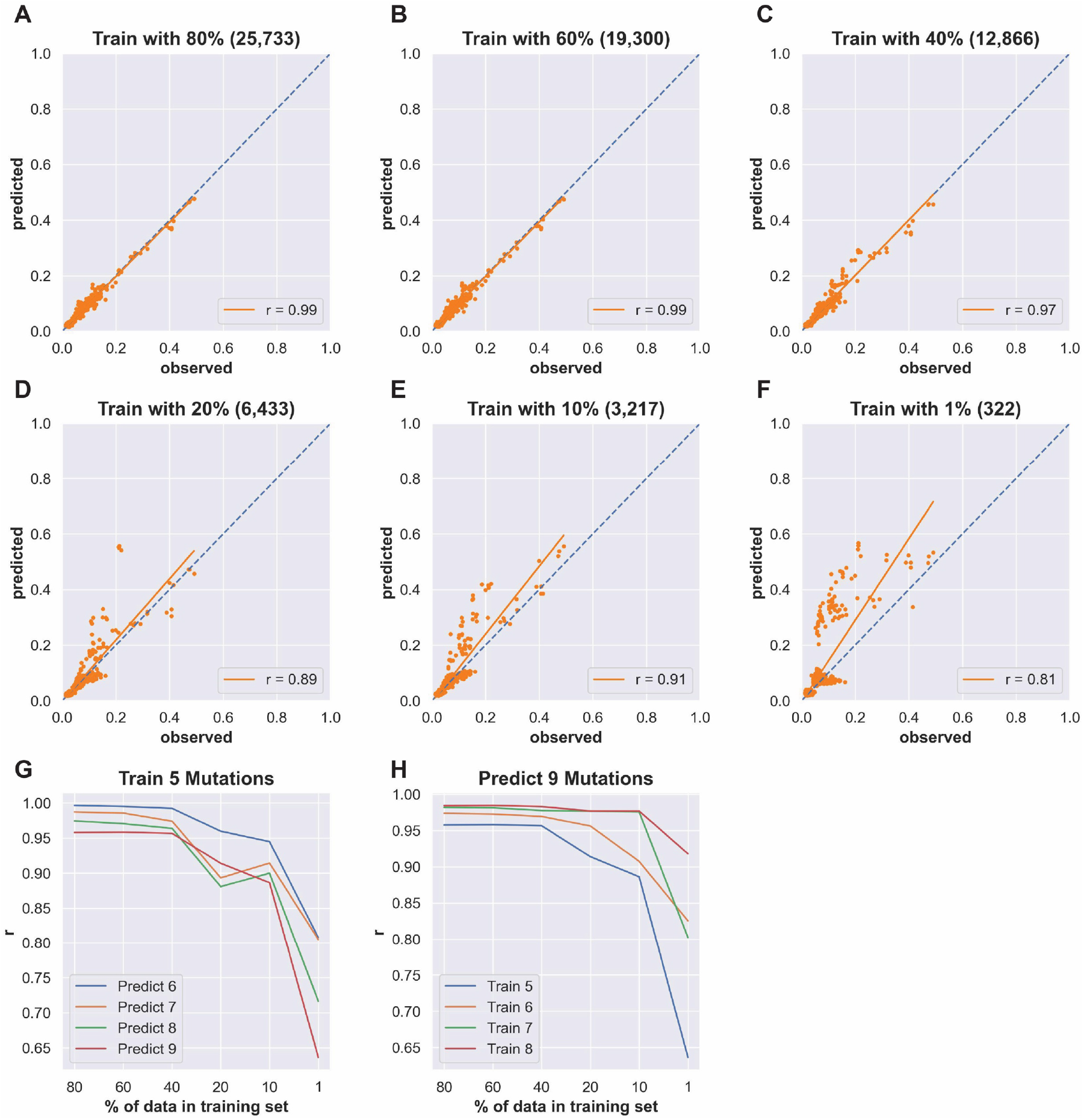
Effects of reducing the number of sequences in the training data. (A-F) Scatter plots of *Predicted* (fraction cleaved from the models) and *Observed* (fraction cleaved from experiments) for models trained with decreasing amounts of sequences with 5 or fewer mutations using the random forest approach and predicting the fraction cleaved of sequences with 7 mutations. The percent of the total sequence used in the training data is indicated in the title of each plot, and the number of unique sequences in the training data is reported in parentheses. Pearson correlation coefficients *r* are indicated as insets. (G) The correlation between predicted and observed for a model trained with decreasing amounts data from sequences with 5 or fewer mutations (“Train 5”) and predicting the activity of sequences with increasing numbers of mutations (Predict 6-9). (H) Predicting the activity of sequences with 9 mutations (“Predict 9 Mutations”) with models trained on different reduced data sets.

While the primary goal was to predict the relative activity of RNA sequences, we wondered if the models might also be useful for predicting structurally important nucleotides. To address this question, we analyzed the “feature importance” in several of our Random Forest models. Feature importance is a method to assign importance to specific input data. Because our data only uses sequence as input, the features in our data are specific nucleotides (A, G, C or U) at specific positions. We found that for the Random Forest models, the most important feature all clustered around the active site of the ribozyme (Supplemental Figures 16 and 17). Further, the CPEB3 ribozyme uses metal ion catalysis and several of the most important features were nucleotides that have been observed coordinated to the active site magnesium ion in the CPEB3 ribozyme, or the analogous nucleotides in the structurally similar HDV ribozyme (Kapral et al., 2014; Skilandat et al., 2016). For example, for all the models trained with some higher order mutants, the most important feature was G1, which positions the cleaved phosphate bond in contact with the catalytic magnesium ion. The second most important feature was G25, which forms a wobble base pair with U20 (Lévesque et al., 2012), another important feature (top 4-6), and this nucleotide pair coordinates the active site magnesium ion through outer sphere contacts. The catalytic nucleotide C57 binds the same catalytic magnesium as the G25:U20 wobble pair, and had a high feature importance similar to U20. Most of the other important features are involved in base pairs that stack or interact with the metal ion coordinating bases. Interestingly, we found that nine of the ten most important features were identical for models trained with only single and double mutants or with increasing amounts of higher-order mutants. However, the G1 and G25 features became increasingly more important as sequences with higher mutational distance were added to the training data. This indicates that the higher-order mutants in the training data helped emphasize structurally critical nucleotides. We conclude that the machine learning models presented identified nucleotides involved in forming the active sites of the CPEB3 ribozyme. Because we did not use structural data to train our models, the results suggest that similar data could identify active sites in RNA molecules with unknown structures.

## Discussion

We have shown that a model trained on ribozyme activity data can accurately predict the self-cleavage activity of sequences with numerous mutations. This approach can be used to guide experiments based on a relatively small set of initial data. Importantly, the approach did not use structural information such as X-ray crystallography or cryo-EM, and used only sequence and activity data, which can be obtained with common molecular biology approaches (in vitro transcription, RT-PCR, and sequencing). In addition, the training data starts with small amounts of synthetic DNA. The comprehensive double mutant data and the phylogenetic derived data each started from a single DNA oligo synthesis that used doped phosphoramidites at the variable positions. Each data set was collected on a single lane of an Illumina sequencer. The approach presented in this paper is therefore accessible, rapid and inexpensive as compared to approaches that use structural data to train their models.

Sequence conservation of naturally occurring RNA molecules has been another useful data type for training models to predict RNA structure from sequence (De Leonardis et al., 2015; Weinreb et al., 2016). This approach is based on the observation that nucleotide positions that form a base pair often show co-evolutionary patterns of sequence conservation. In some cases, this co-evolutionary data has been combined with thermodynamic predictions or structural data from chemical probing, such as SHAPE experiments (Calonaci et al., 2020). Numerous ribozymes, aptamers and aptazymes have been discovered through in vitro evolution experiments and conservation data is not available unless sequencing experiments were applied during the selection process. Our approach could be used to expand functional information of non-natural RNA molecules which could then be used to guide structure prediction of these molecules in a way similar to how naturally occurring sequence conservation has been used. In addition, sequence conservation does not necessarily predict relative activity. For example, while the CPEB3 ribozyme is highly conserved in nature, not all of the sequence are equally proficient at catalyzing self-cleavage (Chadalavada et al., 2010; Bendixsen et al., 2021). Our approach using machine learning from experimentally derived data may prove useful for guiding experiments with non-natural RNA molecules discovered through in vitro selection or SELEX-like approaches. However, adopting this machine learning approach will require that each experimenter acquire specific data for their system necessary to train and test sequences with the functions they are investigating.

With future work, it may be possible to produce more general models of ribozyme activity. For example, a model trained on data sets from several different self-cleaving ribozymes with different nucleotide lengths might learn to predict the activity of sequences of arbitrary length and sequence composition. In fact, recent advances in RNA structure prediction have used the crystal structures of several different self-cleaving ribozymes as training data to develop predictive modes that achieve near-atomic level resolution of arbitrary sequences (Townshend et al., 2021). Alternatively, models trained on ribozymes with different activities beyond self-cleavage might be able to classify sequences as ribozymes of various functions. There has been some success with generating general models for predicting protein functions. The latent features identified by deep generative models of protein sequences are being used to better understand the complex, higher-order amino acid interactions necessary to achieve a functional protein structure (Riesselman et al., 2018; Detlefsen et al., 2022). We hypothesize that latent features could aid in the identification of generalized parameters that govern the epistatic interactions of higher-order mutants of RNA sequences as well. We hope that the accuracy and accessibility of the approach presented here will inspire others to carry out similar experiments and initiate the data sharing that will be needed to develop more general models, similar to what is being accomplished for protein functional predictions (Biswas et al., 2021).

One challenge to our predictive models appears to be the low frequency of active sequences at higher mutational distances. In our phylogenetically derived data the vast majority of sequences have very low activity (Figure 1D), and the probability of finding sequence with high fraction cleaved decreases with the number of mutations relative to wild-type. As a consequence, models trained on lower-order mutant variants tend to overestimate the activity of sequences at higher mutational distances. It has been previously observed that experimental RNA fitness landscapes are dominated by *negative epistasis*, which means that mutations in combination tend to have lower fitness than would be expected from the additive effects of individual mutations (Bendixsen et al., 2017). The overestimation of fraction cleaved at higher mutational distances suggests that our models have a difficult time learning to predict negative epistasis. It has been previously observed that mutations with “neutral” or “beneficial” effects on protein function often have destabilizing effects on protein structure (Soskine and Tawfik, 2010). We postulate that the same effect is causing negative epistasis in the RNA data. This suggests that additional information, such as measurements or estimates of thermodynamic stability of helices, might be necessary for increasing accuracy at even higher distances beyond those offered by this data set (Groher et al., 2019; Yamagami et al., 2019). For example, we have recently demonstrated that our sequencing based approach to measuring ribozyme activity can be extended to include magnesium titrations in order to evaluate RNA folding/stability (Peri et al., 2022). In the future, combining structural and functional information might be the best approach to accurately design RNA molecules with desired functional properties.

## Materials and Methods

### Ribozyme activity data

Ribozyme activity was determined as previously described (Bendixsen et al., 2021). Briefly, DNA templates were synthesized with the promoter for T7 RNA polymerase to enable in vitro transcription. Templates were synthesized with mixtures of phosphoramidites at variable positions. For the comprehensive double-mutant data set, templates were synthesized with 97% wild-type nucleotides and 1% each of the other three nucleotides. For the phylogenetic derived data set, the template was synthesized with an equal mixture of the naturally occurring nucleotides that were found at 13 positions that varied across 99 mammalian genomes. During in vitro transcription, RNA molecules self-cleaved at different rates. The reaction was stopped at 30 minutes, and the RNA was concentrated and reverse transcribed with a 5’-RACE protocol that appends a new primer site to the cDNA of both cleaved and uncleaved RNA (SMARTScribe, Takara). The cDNA was PCR amplified with primers that add the adaptors for Illumina sequencing. This procedure was done in triplicate with unique dual-indexes for each replicate. DNA was combined equimolar and sent for sequencing (GC3F, University of Oregon). Sequencing was performed on a single lane of a HiSeq 4000 using paired-end 150 reads.

### Ribozyme activity from sequence data

FastQ sequencing data were analyzed using custom Julia and Python scripts. Briefly, the scripts identified the reverse transcription primer binding site at the 3’-end to determine nucleotide positions and then determined if the sequence was cleaved or uncleaved by the absence or presence of the 5’-upstream sequence. For the single and double mutants, all possible sequences were generated and stored in a list, and reads that matched the list elements were counted and cleaved or uncleaved was determined by the presence or absence of the 5’-upstream sequence. For the phylogenetically derived data, nucleotide identities were determined at the expected 13 variable positions by counting the string character position from the fixed regions. Sequencing reads were discarded if they contained unexpected mutations in the primer binding site, the uncleaved portion, or the ribozyme sequence. For each unique genotype in the library the number of cleaved and uncleaved sequences were counted and ribozyme activity (fraction cleaved) was calculated as *fraction cleaved* = counts_cleaved_/(counts_cleaved_ + counts_uncleaved_).

### Machine Learning

Random Forest regression uses an ensemble of decision trees to improve prediction accuracy. Each tree in the ensemble is created by partitioning the sequences within a sample into groups possessing little variation. Each sample is drawn with replacement and the resulting trees are aggregated into forests that best predict the cleavage rates of the sequences. The Random Forest regression was performed using the python package scikit-learn. Each sequence was transformed into a 69 by 4 one-hot encoding representation of the sequence. Each of the four possible nucleotides within the sequence was represented by a vector of length 4 possessing a uniquely located “1” within the vector to signify the nucleotide’s identity. Each sequence in the training set was fit using scikit-learn’s RandomForestRegressor ensemble module. Feature importance was computed via a forest of randomized trees using the *features_importances* function in the module under default settings. Briefly, the relative importance of a feature was determined by the depth of the feature when it was used as a decision node in a tree. Features used at the top of the tree contribute to the final prediction decision of a larger fraction of the input samples. The expected fraction of the samples they contributed to was used as an estimate of the relative importance of the features.

LSTM is a recurrent neural network commonly used for the predictive modeling of written text data, which has sequential dependencies. Here we used an LSTM to compute a set of hidden features given a set of nucleotide sequences. These hidden features are learned by the LSTM in a supervised way for the purpose of relating the nucleotide sequence to the corresponding ribozyme activity (fraction cleaved). The LSTM network has an architecture where each cell *C* outputs the next state h_t_ (1 ≤ t ≤ n) by taking in input from the previous state h_t-1_ and the embedding x_t_ of the current nucleotide in the sequence. The output h_n_ of the last cell of the LSTM is then used as input to a Random Forest regressor to predict the sequence functional activity rate. The LSTM model was built using PyTorch’s open-source machine learning framework. Sequences were trained using an LSTM layer with 32 hidden dimensions and a dropout rate of 0.2. Each sequence was embedded in a 69 by 4 tensor (where 4 is the size of the nucleotide embedding) and then batched in groups of 64 sequences for input to the model. The gradient descent was performed using PyTorch’s built-in Adam optimizer and MSELoss criterion. Twenty-five training epochs were performed on each training set.

### Training and Test Data

The data set containing the fraction cleaved data from the 27,647 phylogenetically derived sequences was binned based on the number of mutations relative to the wild-type ribozyme. For each bin, a portion of the data (20%) was chosen at random and set aside as test data. This resulted in test data sets that were also separated by the number of mutations relative to the wild-type sequence. Training data sets were created from the 80% of data in each mutational bin that was not set aside for testing. Training data sets were created by combining bins at a given number of mutations to all the bins with lower numbers of mutations. Training data included 100% of the single and double mutant data. For reduced training sets were created by randomly sampling different numbers of sequences from the original full training data sets.

## Data Availability

Sequencing reads in FastQ format are available at ENA (PRJEB51631). Sequence and activity data and computer code is available at GitLab (https://gitlab.com/bsu/biocompute-public/ml-ribo-predict.git).

## Author Contributions

JB – involved in conceptualizing the project, managed data, performed computational work for formal analysis and visualization, reviewed and edited the manuscript; JR – involved in conceptualizing the project, performed experiments, managed the project, supervised and facilitated computational work, prepared figures, reviewed and edited the manuscript, JK – involved in conceptualizing the project, performed computational work for formal analysis and validation; AT– was involved in conceptualizing the project, helped with experimental validation, supervised and facilitated computational work; ES – Involved in conceptualizing the project, supervised computational work, reviewed and edited the manuscript; FS – Involved in conceptualizing the project, supervised computational work, reviewed and edited the manuscript; EH – involved in conceptualizing the project, supervised experimental work, supervised computational work, wrote the original draft, and reviewed and edited the manuscript.

## Acknowledgments

We thank Devin Bendixsen for producing the fitness landscape image used in the manuscript and for valuable discussions.

## References

Athavale, S. S., Spicer, B., and Chen, I. A. (2014). Experimental fitness landscapes to understand the molecular evolution of RNA-based life. Current Opinion in Chemical Biology 22, 35–39. doi: 10.1016/j.cbpa.2014.09.008.

Bendixsen, D. P., Østman, B., and Hayden, E. J. (2017). Negative Epistasis in Experimental RNA Fitness Landscapes. J. Mol. Evol. doi: 10.1007/s00239-017-9817-5.

Bendixsen, D. P., Pollock, T. B., Peri, G., and Hayden, E. J. (2021). Experimental resurrection of ancestral mammalian CPEB3 ribozymes reveals deep functional conservation. Mol Biol Evol. doi: 10.1093/molbev/msab074.

Biswas, S., Khimulya, G., Alley, E. C., Esvelt, K. M., and Church, G. M. (2021). Low-N protein engineering with data-efficient deep learning. Nat Methods 18, 389–396. doi: 10.1038/s41592-021-01100-y.

Blanco, C., Janzen, E., Pressman, A., Saha, R., and Chen, I. A. (2019). Molecular Fitness Landscapes from High-Coverage Sequence Profiling. Annu Rev Biophys. doi: 10.1146/annurev-biophys-052118-115333.

Calonaci, N., Jones, A., Cuturello, F., Sattler, M., and Bussi, G. (2020). Machine learning a model for RNA structure prediction. NAR Genomics and Bioinformatics 2, lqaa090. doi: 10.1093/nargab/lqaa090.

Chadalavada, D. M., Gratton, E. A., and Bevilacqua, P. C. (2010). The Human HDV-like CPEB3 Ribozyme Is Intrinsically Fast-Reacting. Biochemistry 49, 5321–5330. doi: 10.1021/bi100434c.

De Leonardis, E., Lutz, B., Ratz, S., Cocco, S., Monasson, R., Schug, A., et al. (2015). Direct-Coupling Analysis of nucleotide coevolution facilitates RNA secondary and tertiary structure prediction. Nucleic Acids Res 43, 10444–10455. doi: 10.1093/nar/gkv932.

Detlefsen, N. S., Hauberg, S., and Boomsma, W. (2022). Learning meaningful representations of protein sequences. Nat Commun 13, 1914. doi: 10.1038/s41467-022-29443-w.

Dykstra, P. B., Kaplan, M., and Smolke, C. D. (2022). Engineering synthetic RNA devices for cell control. Nature Reviews Genetics, 1–14.

Ferré-D’Amaré, A. R., and Scott, W. G. (2010). Small self-cleaving ribozymes. Cold Spring Harbor perspectives in biology 2, a003574.

Ferretti, L., Weinreich, D., Tajima, F., and Achaz, G. (2018). Evolutionary constraints in fitness landscapes. Heredity 121, 466–481. doi: 10.1038/s41437-018-0110-1.

Groher, A.-C., Jager, S., Schneider, C., Groher, F., Hamacher, K., and Suess, B. (2019). Tuning the Performance of Synthetic Riboswitches using Machine Learning. ACS Synth. Biol. 8, 34–44. doi: 10.1021/acssynbio.8b00207.

Groher, F., and Suess, B. (2014). Synthetic riboswitches—a tool comes of age. Biochimica et Biophysica Acta (BBA)-Gene Regulatory Mechanisms 1839, 964–973.

Jimenez, R. M., Polanco, J. A., and Lupták, A. (2015). Chemistry and Biology of Self-Cleaving Ribozymes. Trends in Biochemical Sciences 40, 648–661. doi: 10.1016/j.tibs.2015.09.001.

Kapral, G. J., Jain, S., Noeske, J., Doudna, J. A., Richardson, D. C., and Richardson, J. S. (2014). New tools provide a second look at HDV ribozyme structure, dynamics and cleavage. Nucleic Acids Research 42, 12833–12846. doi: 10.1093/nar/gku992.

Kobori, S., and Yokobayashi, Y. (2016). High-Throughput Mutational Analysis of a Twister Ribozyme. Angew. Chem. Int. Ed. Engl. 55, 10354–10357. doi: 10.1002/anie.201605470.

Lévesque, D., Reymond, C., and Perreault, J.-P. (2012). Characterization of the Trans Watson-Crick GU Base Pair Located in the Catalytic Core of the Antigenomic HDV Ribozyme. PLoS ONE 7, e40309. doi: 10.1371/journal.pone.0040309.

Li, C., Qian, W., Maclean, C. J., and Zhang, J. (2016). The fitness landscape of a tRNA gene. Science, aae0568. doi: 10.1126/science.aae0568.

Peri, G., Gibard, C., Shults, N. H., Crossin, K., and Hayden, E. J. (2022). Dynamic RNA fitness landscapes of a group I ribozyme during changes to the experimental environment. Molecular Biology and Evolution, msab373. doi: 10.1093/molbev/msab373.

Pressman, A. D., Liu, Z., Janzen, E., Blanco, C., Müller, U. F., Joyce, G. F., et al. (2019). Mapping a Systematic Ribozyme Fitness Landscape Reveals a Frustrated Evolutionary Network for Self-Aminoacylating RNA. J. Am. Chem. Soc. 141, 6213–6223. doi: 10.1021/jacs.8b13298.

Riesselman, A. J., Ingraham, J. B., and Marks, D. S. (2018). Deep generative models of genetic variation capture the effects of mutations. Nat Methods 15, 816–822. doi: 10.1038/s41592-018-0138-4.

Salehi-Ashtiani, K., Lupták, A., Litovchick, A., and Szostak, J. W. (2006). A Genomewide Search for Ribozymes Reveals an HDV-Like Sequence in the Human CPEB3 Gene. Science 313, 1788–1792. doi: 10.1126/science.1129308.

Schmidt, C. M., and Smolke, C. D. (2021). A convolutional neural network for the prediction and forward design of ribozyme-based gene-control elements. eLife 10, e59697. doi: 10.7554/eLife.59697.

Skilandat, M., Rowinska-Zyrek, M., and Sigel, R. K. O. (2016). Secondary structure confirmation and localization of Mg2+ ions in the mammalian CPEB3 ribozyme. RNA 22, 750–763. doi: 10.1261/rna.053843.115.

Soskine, M., and Tawfik, D. S. (2010). Mutational effects and the evolution of new protein functions. Nat. Rev. Genet 11, 572–582. doi: 10.1038/nrg2808.

Szendro, I. G., Schenk, M. F., Franke, J., Krug, J., and Visser J. A. G. M. de (2013). Quantitative analyses of empirical fitness landscapes. J. Stat. Mech. 2013, P01005. doi: 10.1088/1742-5468/2013/01/P01005.

Townshend, R. J. L., Eismann, S., Watkins, A. M., Rangan, R., Karelina, M., Das, R., et al. (2021). Geometric deep learning of RNA structure. Science 373, 1047–1051. doi: 10.1126/science.abe5650.

Weinreb, C., Riesselman, A., Ingraham, J. B., Gross, T., Sander, C., and Marks, D. S. (2016). 3D RNA and functional interactions from evolutionary couplings. Cell 165, 963–975. doi: 10.1016/j.cell.2016.03.030.

Yamagami, R., Kayedkhordeh, M., Mathews, D. H., and Bevilacqua, P. C. (2019). Design of highly active double-pseudoknotted ribozymes: a combined computational and experimental study. Nucleic Acids Research 47, 29–42. doi: 10.1093/nar/gky1118.

