## Supplementary Materials for "Predicting higher-order mutational effects in an RNA enzyme by machine learning of high-throughput experimental data"

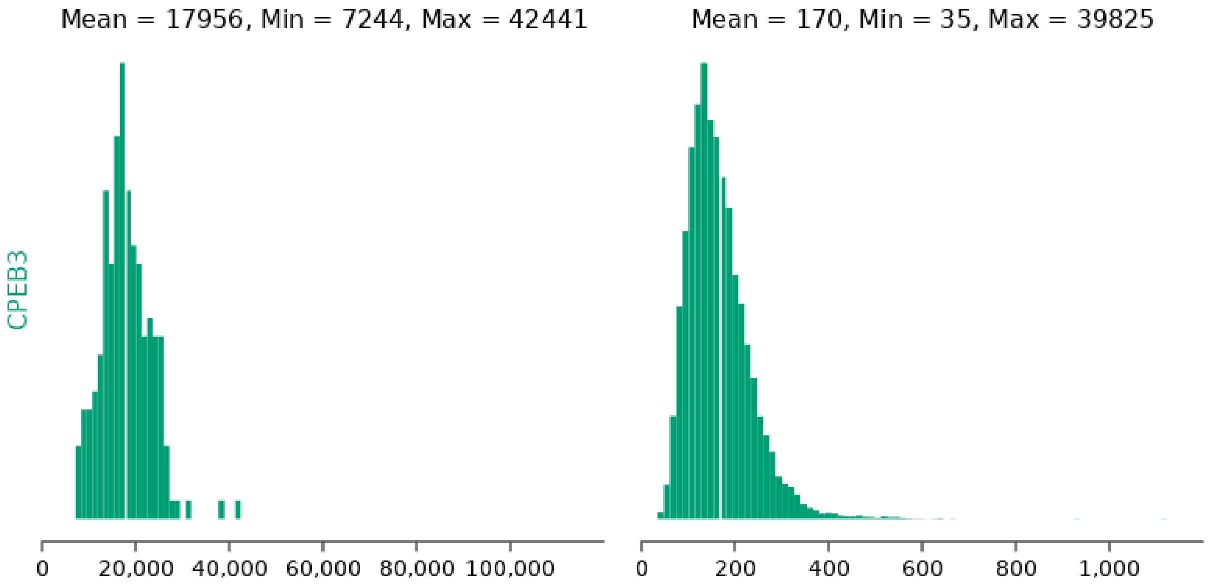


**Supplementary Figure 1.** Histogram of CPEB3 variant counts for single (left) and double (right) mutants. Mean, minimum and maximum values for each distribution are indicated.

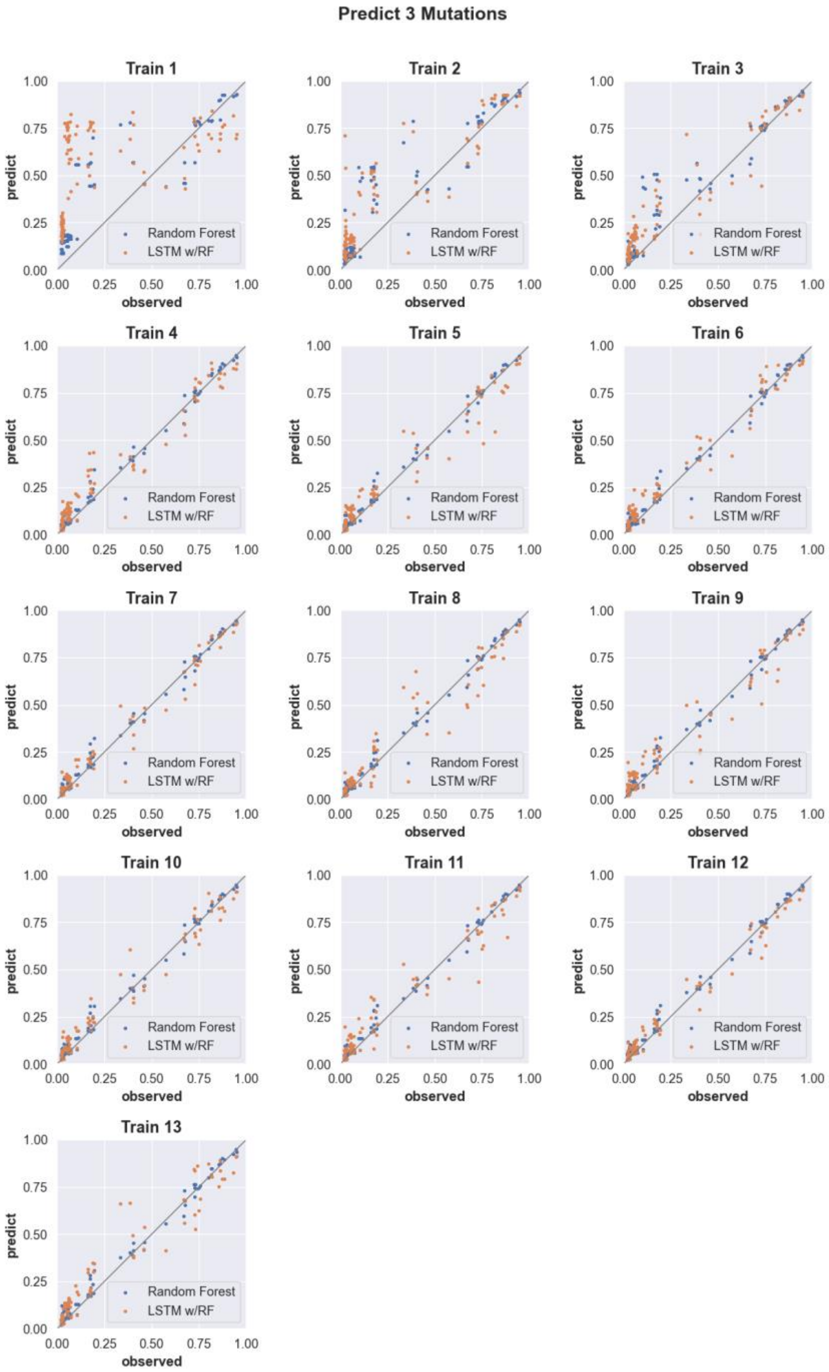


**Supplementary Figure 2. Three-mutation sequence activity predictions.** Scatter plots comparing fraction cleaved values measured from experiments (observed) to those predicted by models (predict) trained by either random forest (blue) or the LSTM approach (orange). Each scatter plot shows the predictions from a different training data set. The training data contained sequences with up to the number of mutations in the title (Train N). For example, 'Train 5' indicates that the model was trained using data for sequences containing 1,2,3,4, and 5 mutations. The line indicates unity, not a fit to the data.

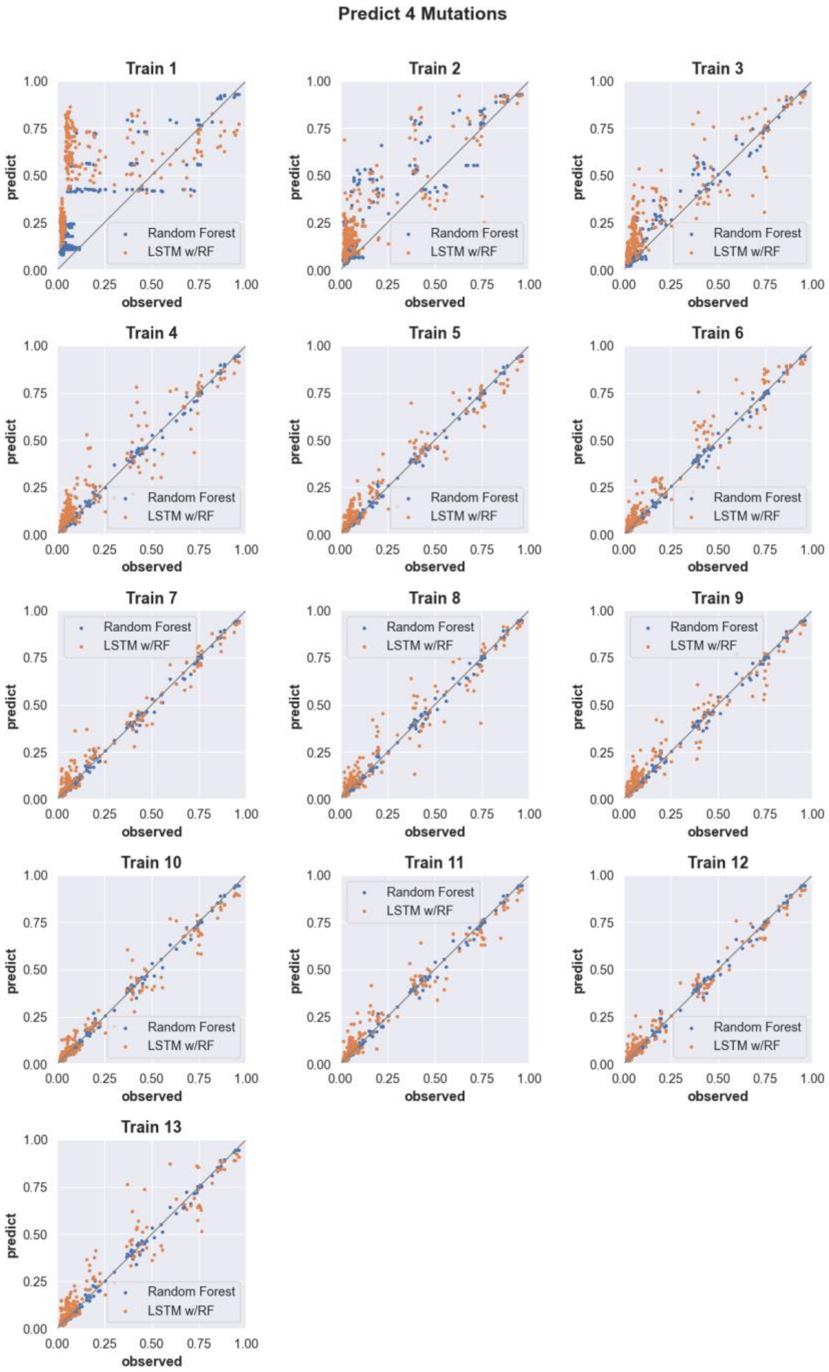


**Supplementary Figure 3. Predicting the activity of sequences with four mutations** (see Supp. Fig. 2 for details).

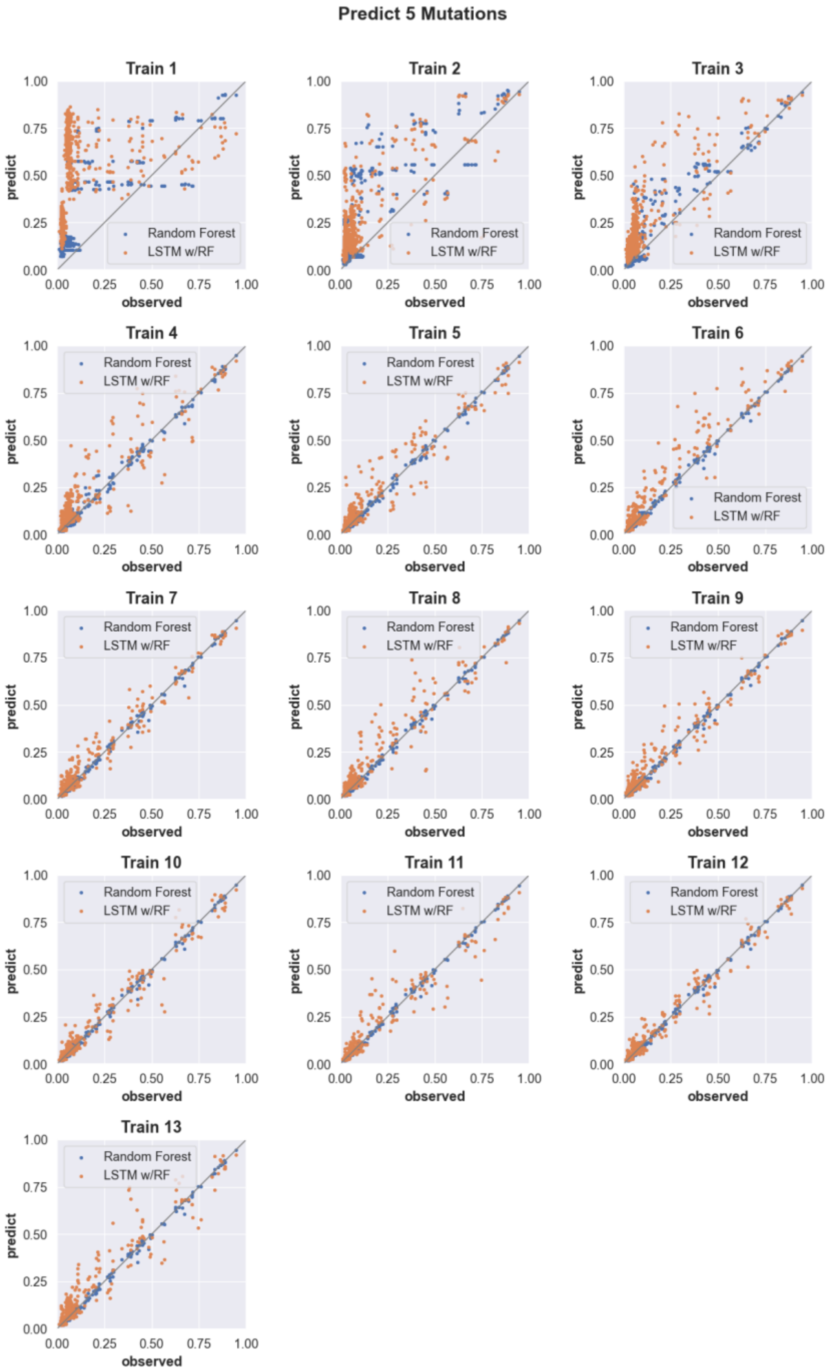


**Supplementary Figure 4. Predicting the activity of sequences with five mutations.** (see Supp. Fig. 2 for details).

.

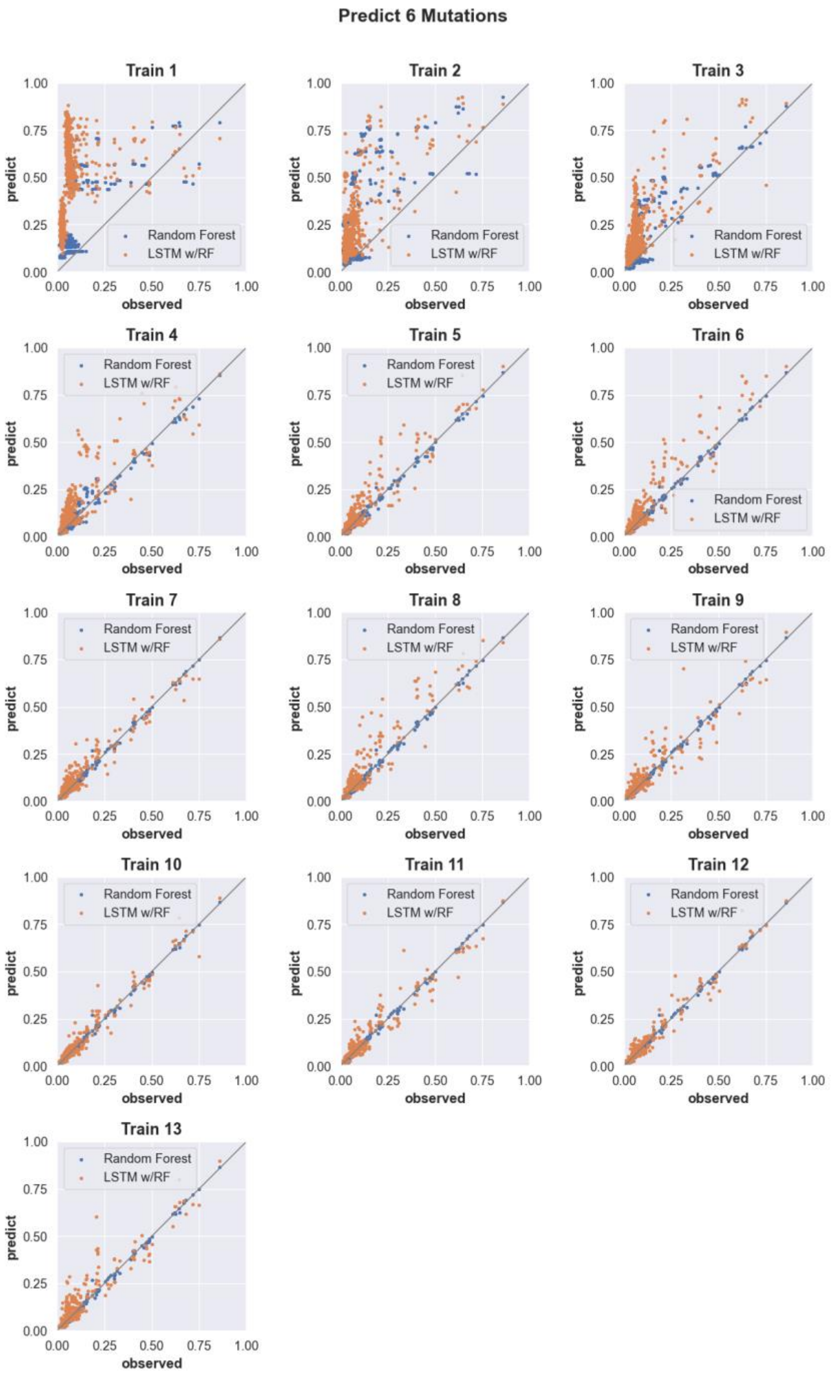


**Supplementary Figure 5. Predicting the activity of sequences with six mutations** (see Supp. Fig. 2 for details).

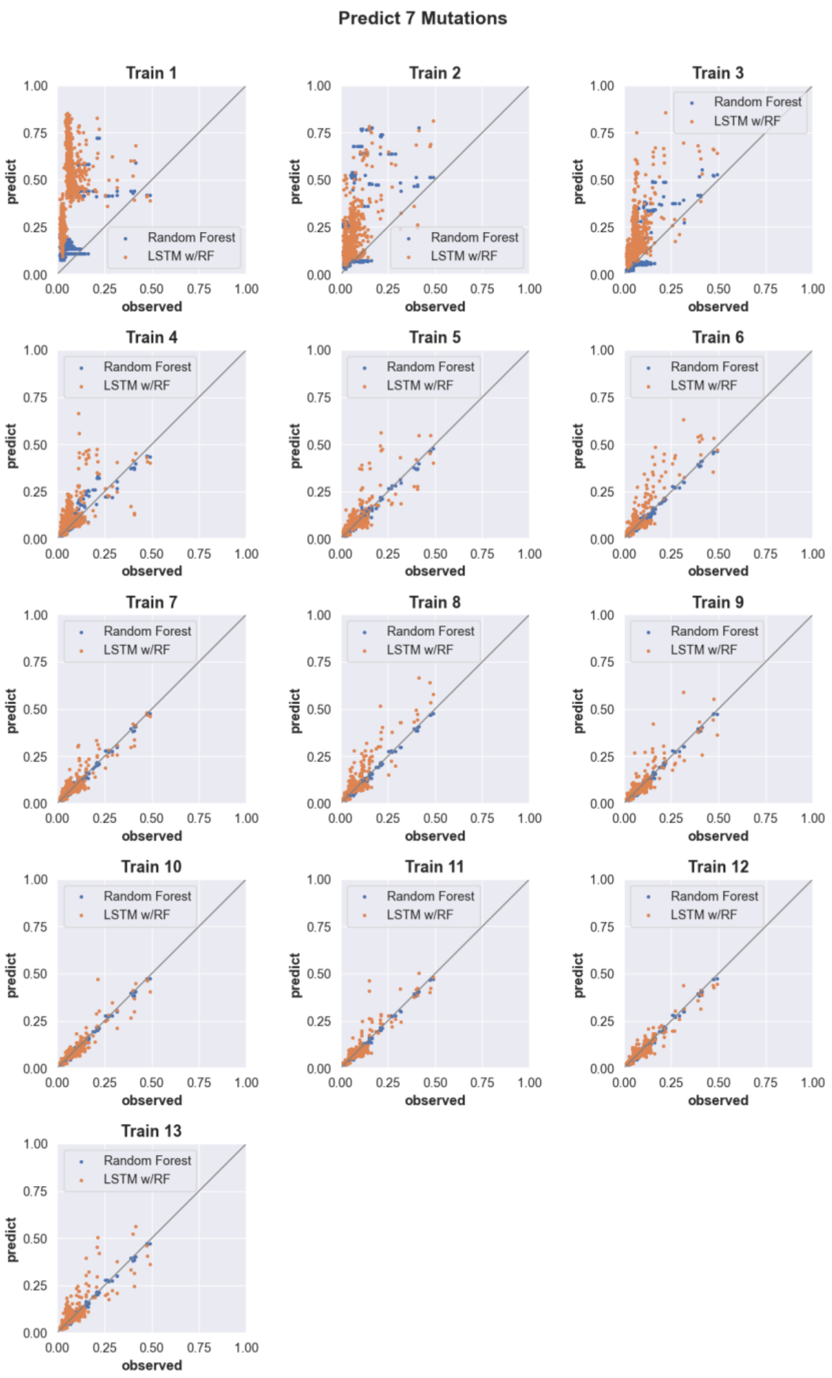


**Supplementary Figure 6. Predicting the activity of sequences with seven mutations** (see Supp. Fig. 2 for details).

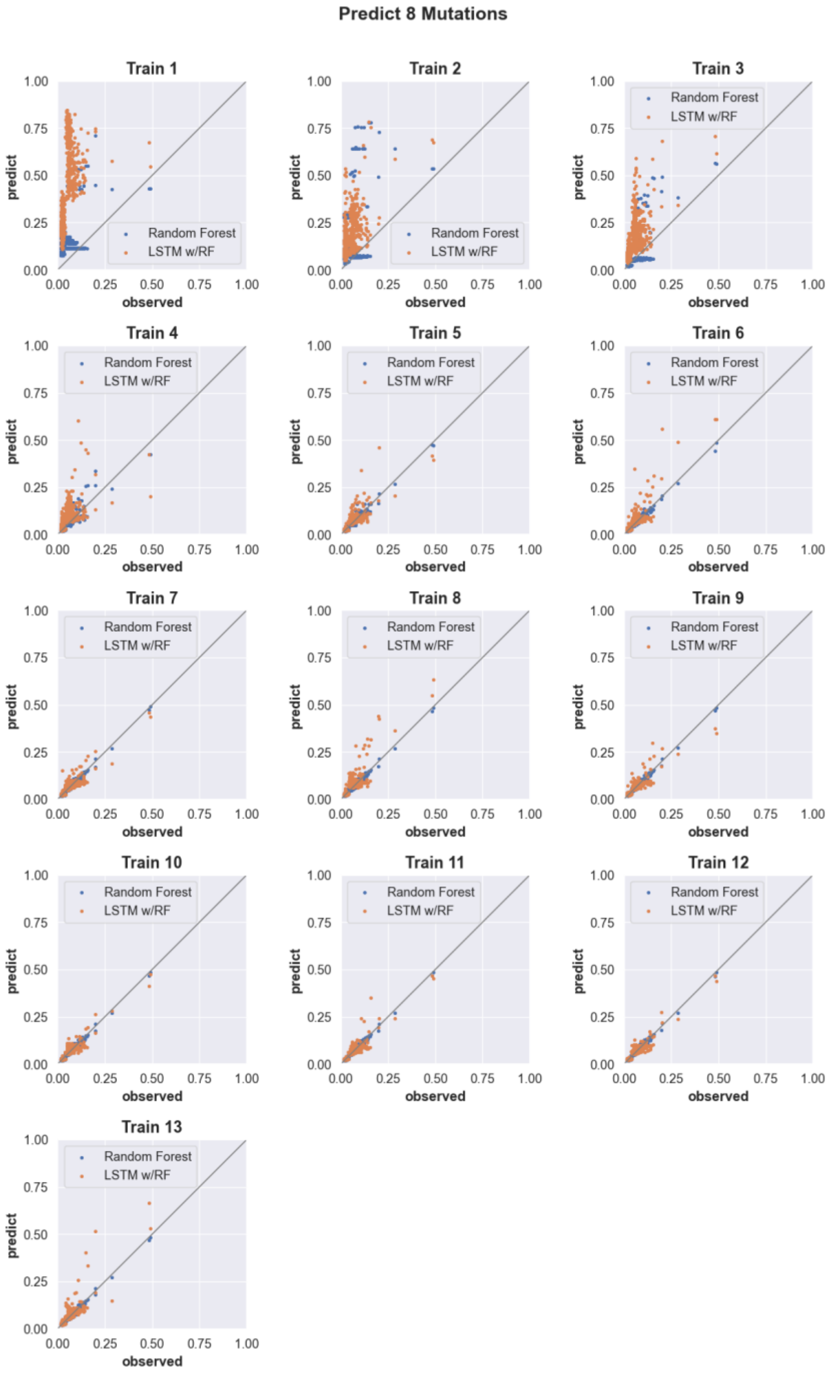


**Supplementary Figure 7. Predicting the activity of sequences with eight mutations** (see Supp. Fig. 2 for details).

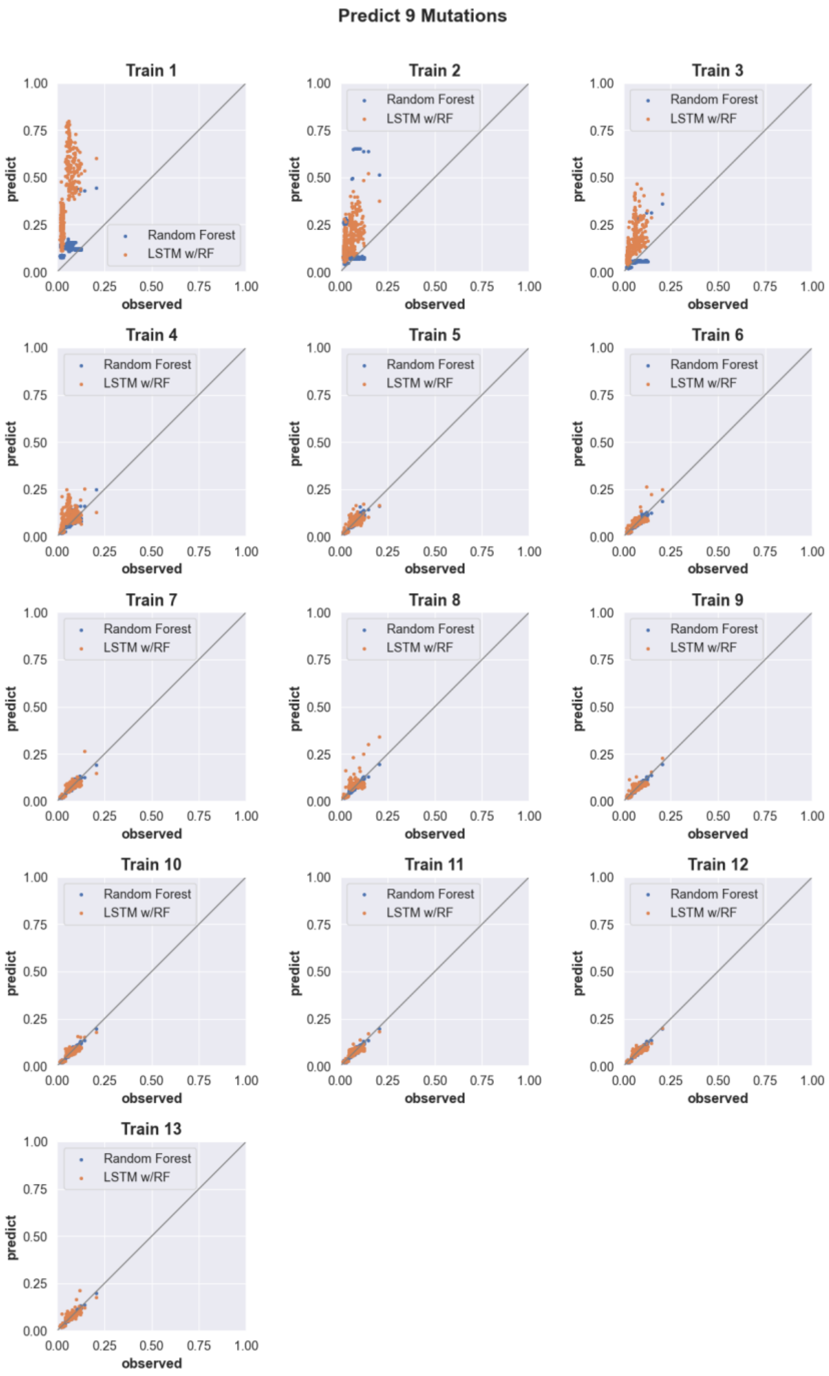


**Supplementary Figure 8. Predicting the activity of sequences with nine mutations** (see Supp. Fig. 2 for details).

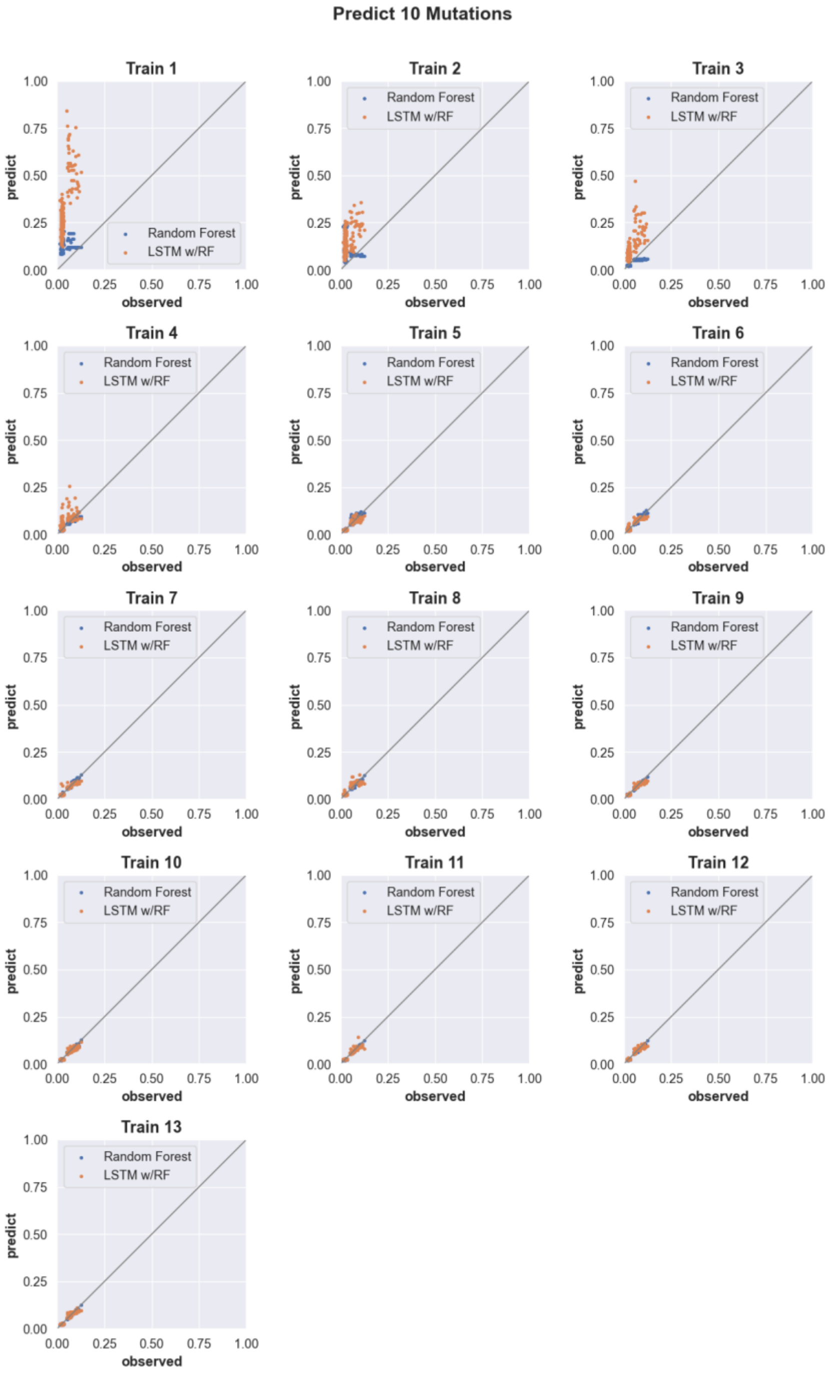


**Supplementary Figure 9. Predicting the activity of sequences with 10 mutations** (see Supp. Fig. 2 for details).

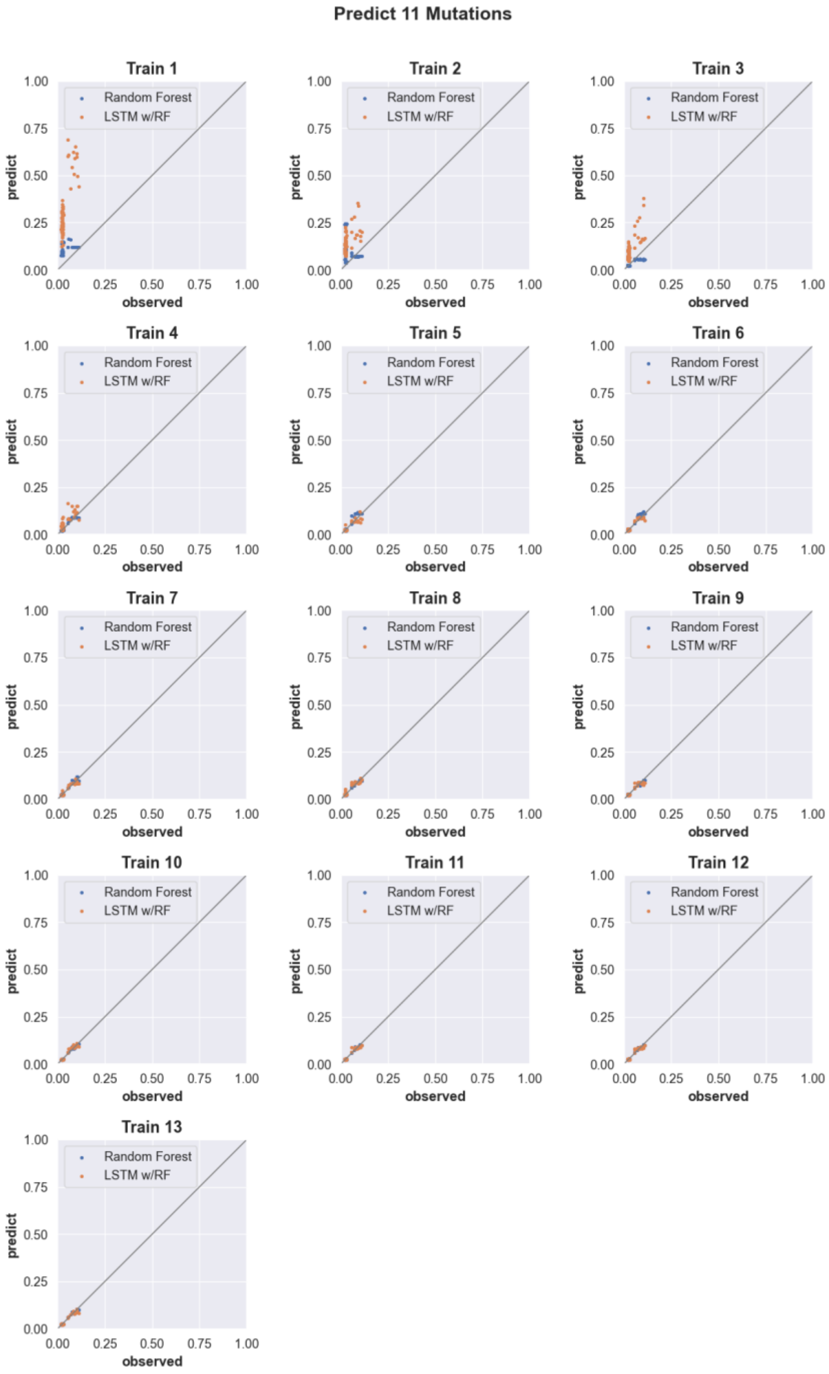


**Supplementary Figure 10. Predicting the activity of sequences with 11 mutations** (see Supp. Fig. 2 for details).

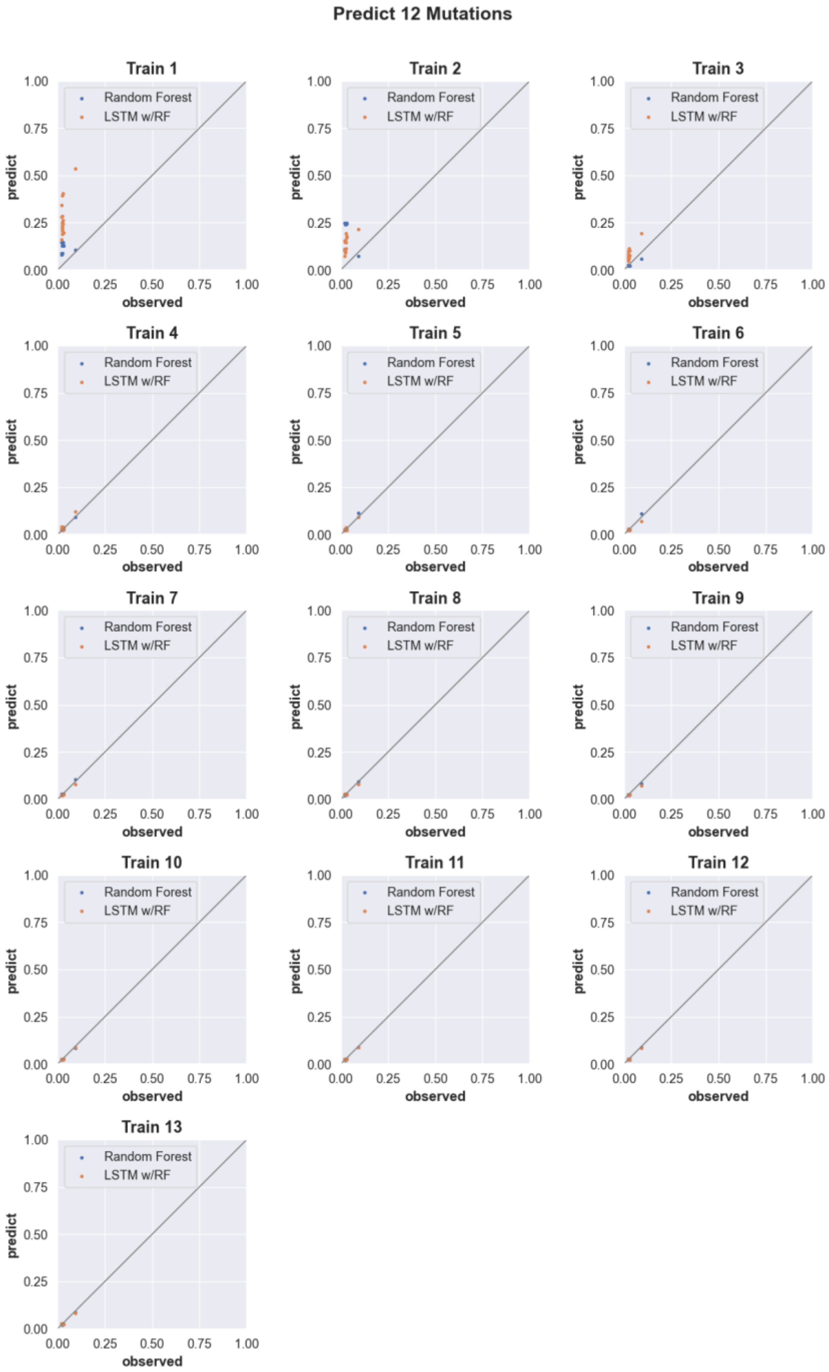


**Supplementary Figure 11. Predicting the activity of sequences with 12 mutations** (see Supp. Fig. 2 for details).

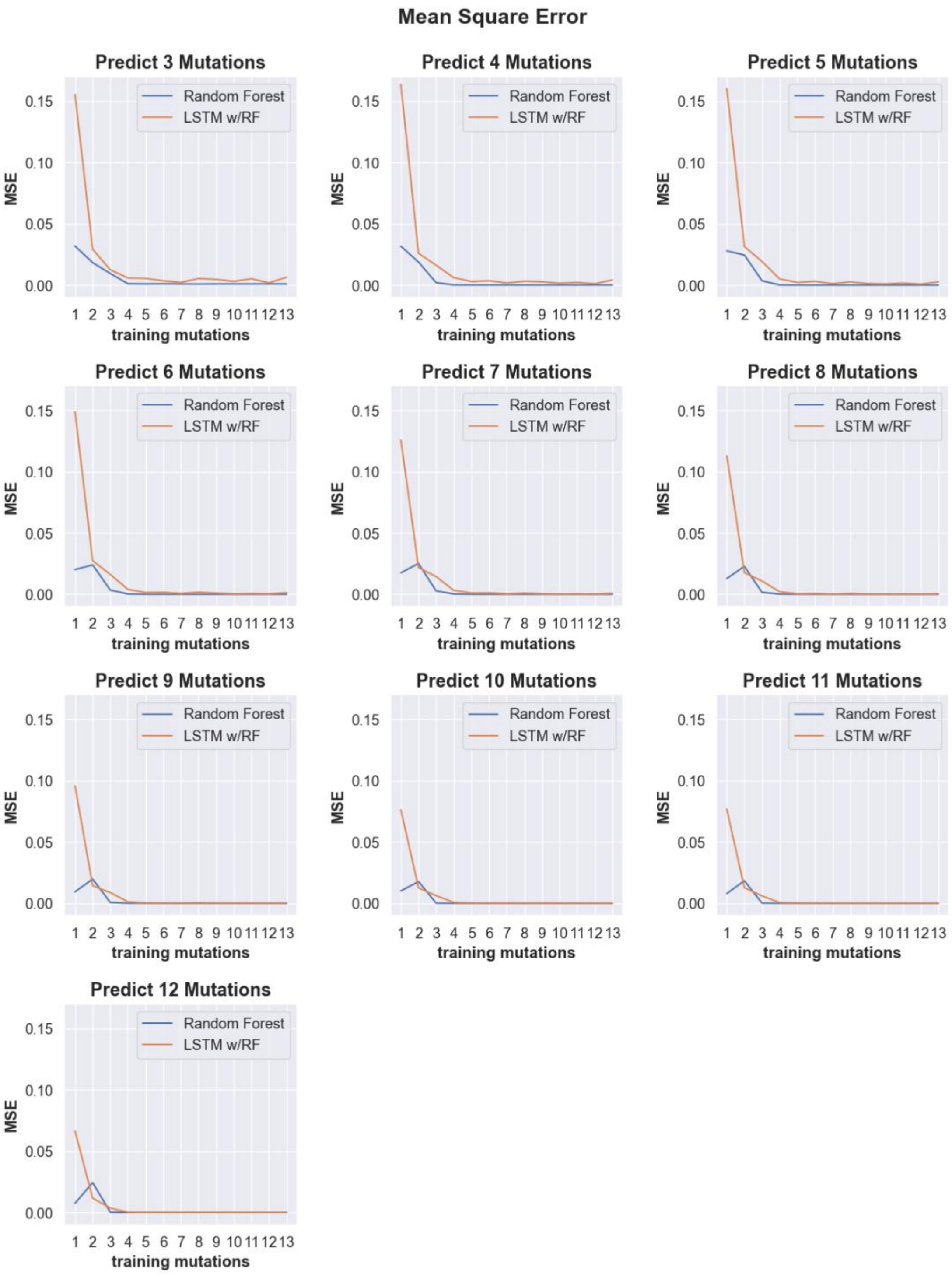


**Supplementary Figure 12.** Line plots showing the mean square error (MSE) of predicted cleavage activity values obtained from random forest (blue) and LSTM with random forest (orange) machine learning models trained on data with incrementally increasing numbers of mutations shown along the x-axis. Each plot shows the MSE for predictions obtained for sequences containing the number of mutations indicated by the plot title.

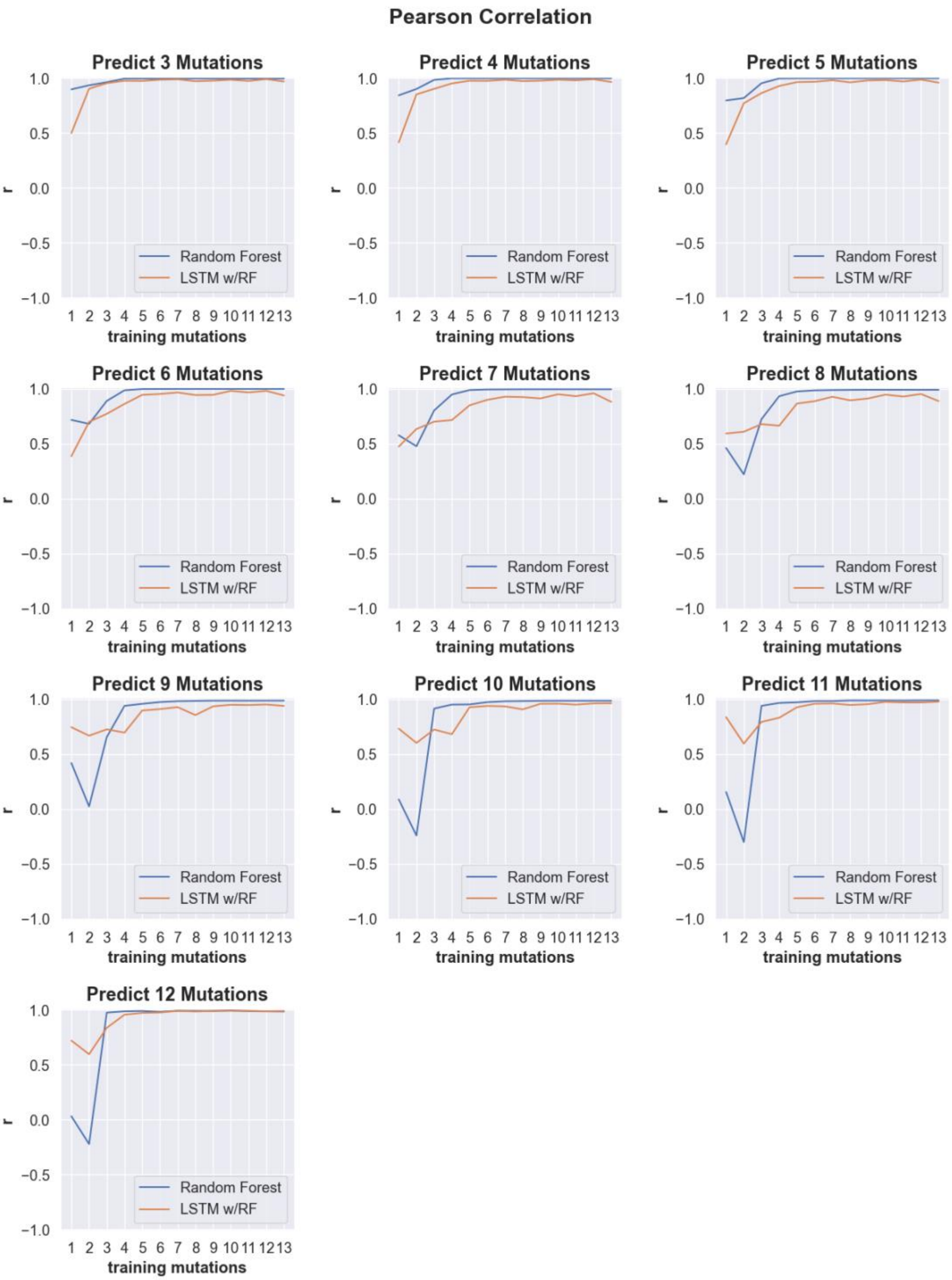


**Supplementary Figure 13.** Line plots showing the Pearson correlation values of predicted cleavage activity obtained from random forest (blue) and LSTM with random forest (orange) machine learning models trained on data with incrementally increasing numbers of mutations shown along the x-axis. Each plot shows the Pearson correlation for predictions obtained for sequences containing the number of mutations indicated by the plot title.

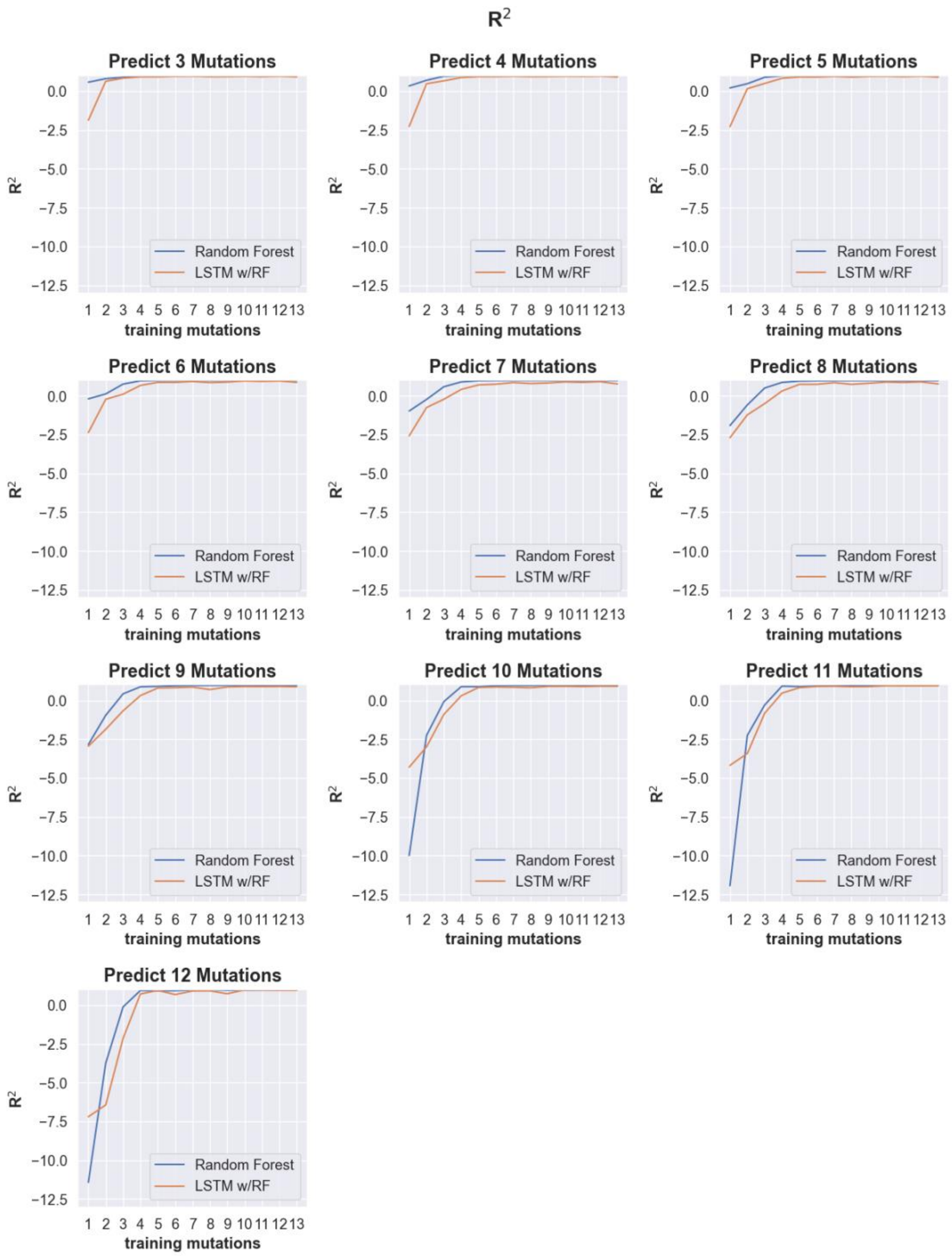


**Supplementary Figure 14.** Line plots showing the $R^2$ values of predicted cleavage activity obtained from random forest (blue) and LSTM with random forest (orange) machine learning models trained on data with incrementally increasing numbers of mutations shown along the x-axis. Each plot shows the $R^2$ for predictions obtained for sequences containing the number of mutations indicated by the plot title.

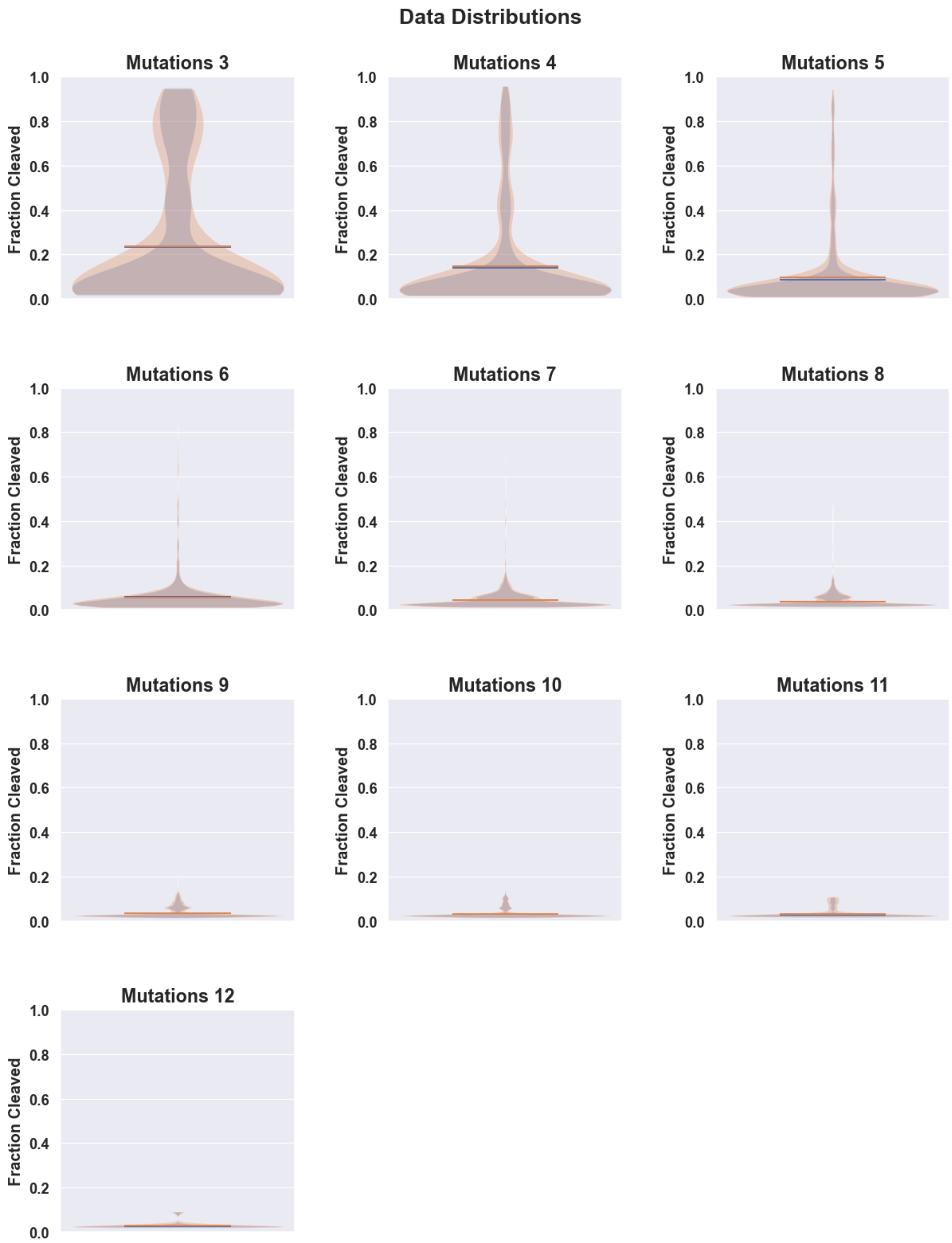


**Supplementary Figure 15.** Violin plots showing the distribution of cleavage rates observed in the test data (orange) and the total data set for a given mutation (blue). The distributions are shown separately for each data set containing increasing numbers of mutations, from 3 to 12.

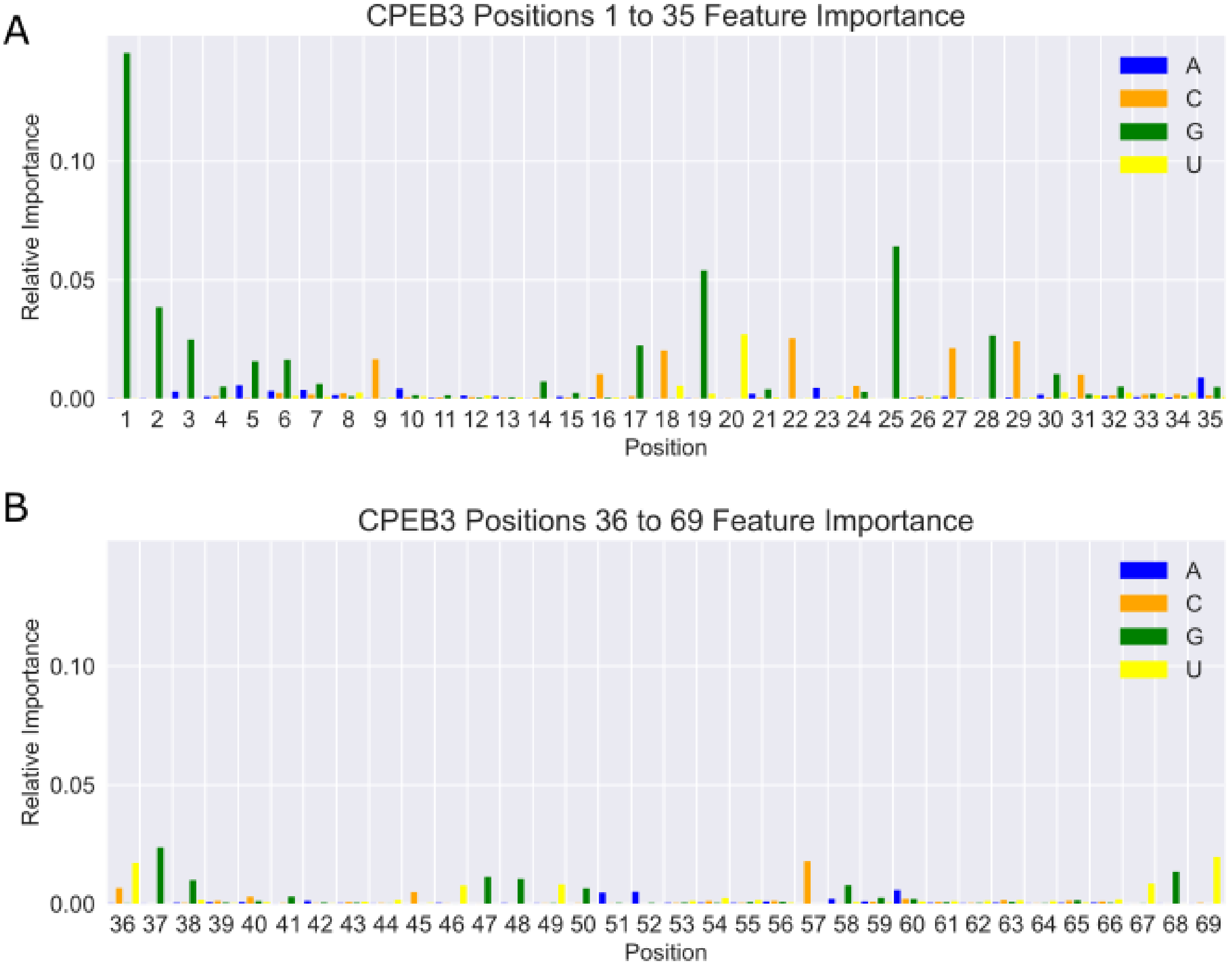


C

Top Ten Feature Importance

| Rank | Train 2 | Train 3 | Train 4 | Train 5 | Train 6 | Train 7 | Train 8 | Train 9 | Train 10 | Train 11 | Train 12 | Train 13 |
|---|---|---|---|---|---|---|---|---|---|---|---|---|
| 1 | G2:0.044 | G1:0.049 | G1:0.084 | G1:0.145 | G1:0.207 | G1:0.251 | G1:0.278 | G1:0.294 | G1:0.301 | G1:0.304 | G1:0.305 | G1:0.305 |
| 2 | G1:0.041 | G2:0.046 | G25:0.046 | G25:0.064 | G25:0.082 | G25:0.093 | G25:0.096 | G25:0.1 | G19:0.096 | G25:0.101 | G25:0.098 | G25:0.096 |
| 3 | G25:0.034 | G25:0.037 | G2:0.043 | G19:0.054 | G19:0.068 | G19:0.08 | G19:0.089 | G19:0.089 | G25:0.095 | G19:0.09 | G19:0.092 | G19:0.095 |
| 4 | T20:0.034 | G28:0.034 | G19:0.042 | G2:0.039 | G2:0.032 | G2:0.028 | G2:0.027 | G2:0.026 | G2:0.026 | G2:0.026 | G2:0.026 | G2:0.026 |
| 5 | G28:0.033 | T20:0.034 | T20:0.032 | T20:0.027 | C29:0.024 | C29:0.025 | C29:0.024 | C29:0.024 | C29:0.023 | C29:0.021 | C29:0.022 | C29:0.02 |
| 6 | C22:0.033 | G19:0.033 | G28:0.032 | G28:0.027 | T20:0.024 | T20:0.021 | T20:0.019 | T20:0.019 | T20:0.018 | T20:0.018 | T20:0.018 | T20:0.018 |
| 7 | G19:0.032 | C22:0.033 | C22:0.03 | C22:0.026 | G28:0.024 | G28:0.021 | G28:0.019 | G28:0.018 | G28:0.018 | G28:0.018 | G28:0.018 | G28:0.018 |
| 8 | G3:0.032 | G3:0.032 | G3:0.029 | G3:0.025 | C22:0.023 | C22:0.02 | C22:0.018 | C22:0.018 | C22:0.017 | C22:0.017 | C22:0.017 | C22:0.017 |
| 9 | G37:0.03 | G37:0.029 | G37:0.027 | C29:0.024 | G3:0.022 | G3:0.019 | G3:0.018 | G3:0.017 | G3:0.017 | G3:0.017 | G3:0.017 | G3:0.017 |
| 10 | T69:0.029 | T69:0.026 | G17:0.024 | G37:0.024 | G37:0.02 | G37:0.018 | T36:0.017 | T36:0.016 | T36:0.016 | T36:0.016 | T36:0.016 | T36:0.015 |
| Total | 0.342 | 0.353 | 0.387 | 0.454 | 0.527 | 0.576 | 0.606 | 0.621 | 0.628 | 0.627 | 0.629 | 0.627 |

**Supplementary Figure 16.** Summary of important features extracted from random forest models. A-B) Bar graphs of feature importance when training with up to five mutations. Each feature represents a specific nucleotide at a specific location, as indicated by the X-axis label (position), and color (nucleotide identity). Positions 1-35 are shown in (A), and positions 36-69 are shown in (B). The height of the bar indicates the relative importance. C) Table ranking the top ten important features extracted from random forest models trained with increasing numbers of mutations. Nucleotides discussed in the main text are highlighted.

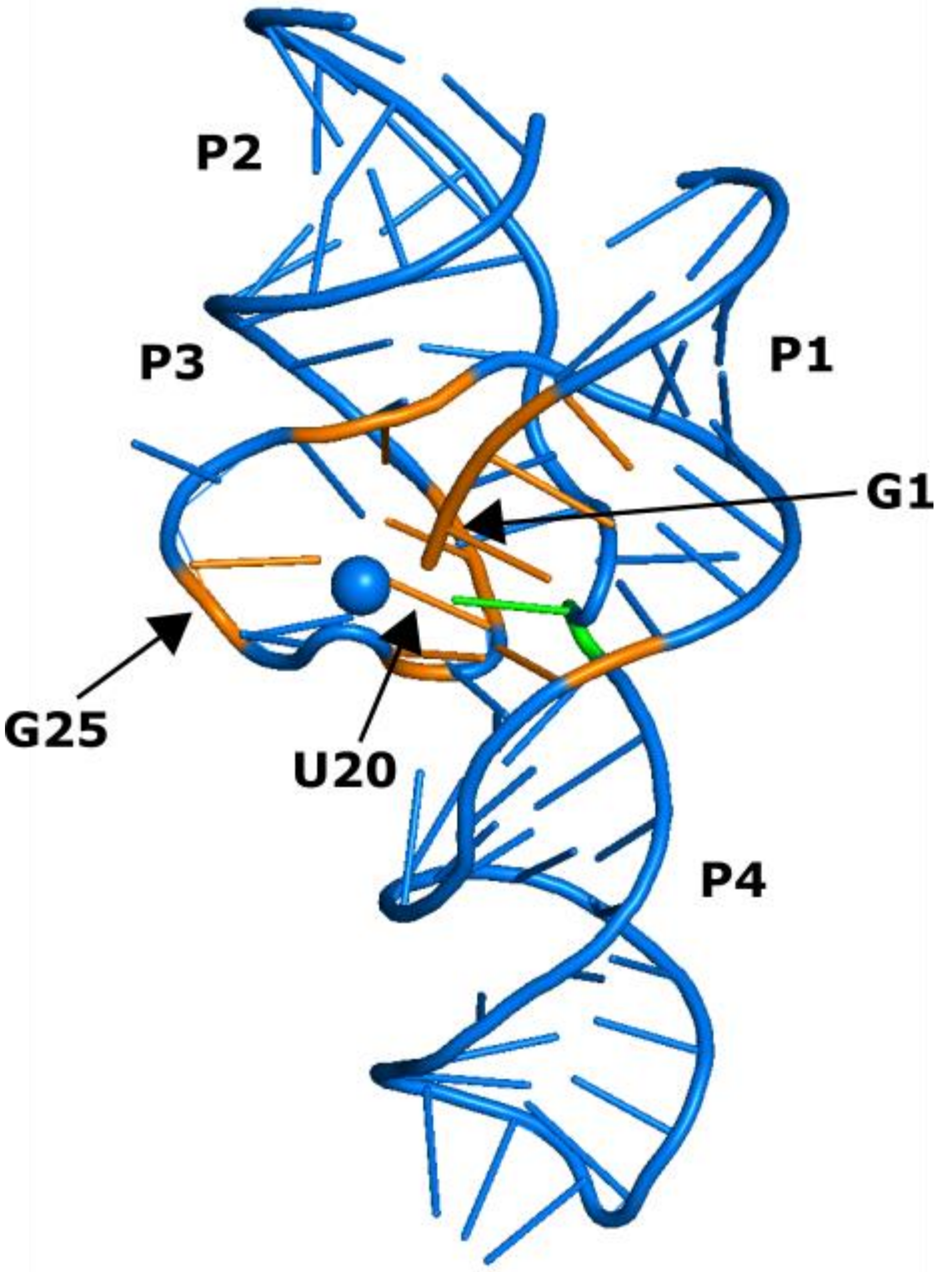


**Supplementary Figure 17.** Crystal structure of an HDV ribozyme (PDB 3NKB) showing the CPEB3 analogous positions representing the top ten important features identified in our random forest models. The feature importance depicted was extracted from the random forest model trained on CPEB3 data including up to 5 mutations. The nucleotides identified as the top ten important features are shaded in orange, the catalytic nucleotide is shaded green (C57/75), and the catalytic $Mg^{2+}$ ion is depicted as a blue sphere.

| Train with % | Pearson | Spearman |
|---|---|---|
| 80% | 0.99 | 0.81 |
| 60% | 0.99 | 0.80 |
| 40% | 0.97 | 0.79 |
| 20% | 0.89 | 0.80 |
| 10% | 0.91 | 0.77 |
| 1% | 0.81 | 0.71 |

**Supplementary Table 1.** Table comparing Pearson and Spearman correlation metrics for reduced training sets containing sequences with up to 5 mutations predicting sequences with 7 mutations. Both Pearson and Spearman correlations show similar, limited reductions in correlation as training set size is reduced.

| Predict Mutations | LSTM w/RF Pearson | RF Pearson | LSTM w/RF Spearman | RF Spearman |
|---|---|---|---|---|
| 3 | 0.9 | 0.93 | 0.80 | 0.78 |
| 4 | 0.85 | 0.9 | 0.60 | 0.81 |
| 5 | 0.77 | 0.82 | 0.52 | 0.82 |
| 6 | 0.7 | 0.68 | 0.51 | 0.74 |
| 7 | 0.63 | 0.48 | 0.44 | 0.7 |
| 8 | 0.61 | 0.22 | 0.46 | 0.64 |
| 9 | 0.67 | 0.02 | 0.50 | 0.63 |
| 10 | 0.6 | −0.24 | 0.45 | 0.51 |
| 11 | 0.6 | −0.3 | 0.48 | 0.37 |

**Supplementary Table 2.** Table comparing Pearson and Spearman correlation metrics for training set containing sequences with up to 2 mutations predicting sequences with 3 to 11 mutations using the LSTM with Random Forest and the Random Forest models. Both Pearson and Spearman correlations show similar reductions in correlation as predictive distance grows.
